# Severe fire regimes decrease resilience of ectothermic populations

**DOI:** 10.1101/2023.06.25.546448

**Authors:** Heitor Campos de Sousa, Adriana Malvasio, Guarino Rinaldi Colli, Roberto Salguero-Gómez

**Affiliations:** Universidade Federal do Tocantins – UFT, Quadra 109 Norte Av. NS-15, ALCNO-14, Plano Diretor Norte, CEP: 77001-90, Palmas, Tocantins, Brasil; Department of Zoology, University of Oxford, 11a Mansfield Rd, OX1 3SZ, UK; Evolutionary Demography Laboratory, Max Planck Institute for Demographic Research, Konrad Zuße straße 1, DE 18057, Germany; Universidade de Brasília – UnB, Departamento de Zoologia, Instituto de Ciências Biológicas, Avenida L4 Norte, Asa orte, CEP: 70910-900, Brasília, Distrito Federal, Brasil

**Keywords:** Demography, Disturbance, Ecophysiology, Environmental stochasticity, Environmental change, Integral Projection Model, Life history, Reptile, Transient dynamics

## Abstract

1. Understanding populations’ responses to environmental change is crucial for mitigating human-induced disturbances.
2. Here, we test hypotheses regarding how three essential components of demographic resilience (resistance, compensation, and recovery) co-vary along the distinct life histories of three lizard species exposed to variable, prescribed fire regimes.
3. Using a Bayesian hierarchical framework, we estimate vital rates (survival, growth, and reproduction) with 14 years of monthly individual-level data and mark-recapture models to parameterize stochastic Integral Projection Models from five sites in Brazilian savannas, each historically subjected to different fire regimes. With these models, we investigate how weather, microclimate, and ecophysiological traits of each species influence their vital rates, emergent life history traits, and demographic resilience components in varying fire regimes.
4. Overall, weather and microclimate are better predictors of the species’ vital rates, rather than their ecophysiological traits. Our findings reveal that severe fire regimes increase populations’ resistance, but decrease compensation or recovery abilities. Instead, populations have higher compensatory and recovery abilities at intermediate degrees of fire severity. Additionally, we identify generation time and reproductive output as predictors of resilience trends across fire regimes and climate. Our analyses demonstrate that the probability and quantity of monthly reproduction are the proximal drivers of demographic resilience across the three species.
5. Our findings suggest that populations surpass a tipping point in severe fire regimes and achieve an alternative stable state to persist. Thus, higher heterogeneity in fire regimes can increase the reproductive aspects and resilience of different populations and avoid high-severity regimes that homogenize the environment. Despite being more resistant, species with long generation times and low reproductive output take longer to recover and cannot compensate as much as species with faster paces of life. We emphasize how reproductive constraints, such as viviparity and fixed clutch sizes, impact the ability of ectothermic populations to benefit and recover from disturbances, underscoring their relevance in conservation assessments.

## 1 INTRODUCTION

Environmental change is causing a fast decline in biodiversity worldwide (Newbold *et al*. 2020). The most impactful drivers of this loss include habitat degradation (Gonçalves-Souza, Verburg & Dobrovolski 2020) and the drastic modification of disturbance regimes (Hale *et al*. 2016), such as fires (Bowman *et al*. 2020). Understanding how species may respond to these impacts is at the core of ecology (Sutherland *et al*. 2013) and conservation biology (Sutherland *et al*. 2009). Ecological resilience research in the last decades has obtained variable success regarding how species traits can predict their responses to disturbances (Halpern 1989; Hu, Doherty & Jessop 2020). Indeed, some organismal traits confer species with higher tolerance or susceptibility to perturbations. Examples include mobility (Merrien *et al*. 2022), robustness (Dantas *et al*. 2013), rapid growth (Hoffmann *et al*. 2012), thermal performance (Hu, Doherty & Jessop 2020), and reproductive output (Capdevila *et al*. 2022). Therefore, deciphering which traits confer species the ability to resist and recover is critical to understanding and predicting how populations will respond to disturbances in the Anthropocene (Carmona *et al*. 2021).

Ecological resilience, the ability of an ecological system to resist and recover, or move to a new stable equilibrium from disturbances (Hodgson, McDonald & Hosken 2015), has been historically explored at the community and ecosystem levels (Kefi *et al*. 2019). Recently, though, key breakthroughs have been made in unraveling the mechanisms behind population responses to disturbances (Capdevila *et al*. 2020). Natural populations may respond to disturbances by avoiding decreases in size (resistance) or even by increasing in size (compensation). Following a transient period caused by the disturbance, populations may recover from said disturbance by transitioning back to the previous or a new stable stationary structure, at which point the composition of stages (*e.g*., age, size classes) in the population remains relatively constant over time. Thus, we can quantify the full repertoire of resilience responses to a disturbance by a natural population with three essential components: (1) resistance, (2) compensation, and (3) recovery time. This novel demographic resilience framework has the advantage of quantifying resilience based on the species’ vital rates (*i.e.*, survival, growth, and reproduction), allowing for inter- and intra-specific comparisons, and providing a mechanistic understanding of the processes and trade-offs involved in species’ responses to disturbances (Capdevila *et al*. 2020).

Life history traits, key moments in an organism’s life cycle (Stearns 1976), have gained attention for their predictive potential in organismal responses (Allen, Street & Capellini 2017). These traits, including generation time and net reproductive output, are governed by survival, growth, and reproduction schedules and are subject to strong selection for optimization (Stearns 2000). As these vital rates involve trade-offs, so do the emerging life history traits (Stearns 1976). Understanding the combination of these traits, known as life history strategies, is valuable in predicting population responses to environmental changes and disturbances (Capdevila *et al*. 2021a). For instance, recent research has shown that plant species with high reproductive output exhibit greater resistance and compensation to disturbances (Iles *et al*. 2015), while longer-lived animals have higher resistance and longer recovery times compared to fast-living species (Capdevila *et al*. 2022). Despite these key findings, their generalization across the Tree of Life is challenging because organisms present diverse life history traits (Salguero-Gómez *et al*. 2016a; Healy *et al*. 2019) and because quantifying trade-offs in natural populations under control conditions is difficult (Metcalf 2016), let alone in a context of disturbance regimes (Lytle 2001). Still, linking mechanisms regulating vital rates to population-level responses to disturbances is crucial for understanding how these trade-offs shape evolutionary processes.

Fire is a major disturbance that shapes ecosystems worldwide (Pausas & Parr 2018). Fire consumes biomass, recycles nutrients, and triggers the renewal of vegetation, leading to a burst of productivity and providing establishment opportunities for new individuals and species (Pausas & Bond 2020). Despite regional heterogeneity in fire activity, the average fire season had globally increased by 18.7% in the last three decades, and in many regions, fire frequency, intensity, and spatial extent have also increased, presumably because of climate change and human management (Bowman *et al*. 2020). In fire-prone ecosystems, species typically develop adaptations to resist and recover from frequent fires (Keeley *et al*. 2011). Conversely, other species have evolved to take advantage of these disturbances. Examples include fire-adapted plants that only resprout (Pausas, Keeley & Schwilk 2017) or whose seed dormancy only breaks (Paniw *et al*. 2016) after a fire, thus benefiting from the open space in temporarily less competitive environments. Historically, though, fire ecology research has primarily focused on plant responses, leaving the mechanisms that confer animals with resilience to fire poorly understood (but see Pausas, Keeley & Schwilk 2017).

To fill this knowledge gap, we investigate how the demographic resilience components (resistance, compensation, and recovery time) co-vary with generation time, reproductive output, and their underlying vital rates. To do so, we parameterize hierarchical Bayesian Integral Projection Models (IPMs) with a 14-years of monthly mark-recapture data from 3,538 individuals of three lizard species (*Copeoglossum nigropunctatum, Micrablepharus atticolus*, and *Tropidurus itambere*) from the Brazilian Cerrado savannas (Oliveira & Marquis 2002), with multiple populations exposed to prescribed fires with varying frequencies and timings. Our three species have rather distinct life-history strategies but co-exist in all of the study sites, thus creating a perfect *in situ* experimental laboratory to explore how their life history traits and resilience components change in response to fire regimes. *Copeoglossum nigropunctatum* (Scincidae) is a medium-sized (up to 150 mm snout-vent length; SVL) viviparous skink found in Amazonia and Cerrado (Vitt & Blackburn 1991; Vitt, Zani & Lima 1997); its females can carry up to nine embryos for 10-12 months (Vitt & Blackburn 1991). *Micrablepharus atticolus* (Gymnophthalmidae) is a small-sized (up to 50 mm SVL) Cerrado endemic with a fixed clutch of 1-2 eggs (Vitt & Caldwell 1993; Rodrigues 1996), but capable of producing multiple clutches (Vieira *et al*. 2000). *Tropidurus itambere* (Tropiduridae) is a medium-sized (up to 100 mm SVL) species from central Cerrado that lays multiple clutches of 5-8 eggs per season (Van Sluys & Henderson 1993). All three species have generalist diets, primarily feeding on small invertebrates, with ants being the main prey for *T. itambere* (Vitt & Blackburn 1991; Van Sluys 1993; Vieira *et al*. 2000). While most lizard species can withstand intense fires (Costa *et al*. 2013), our species respond differently to fire-induced vegetation changes, influenced by microclimate and microhabitat preferences (Costa *et al*. 2021).

Here, we test whether (H1) ecophysiological traits, such as thermal locomotor performance and daily activity, and weather/microclimate shape how underlying vital rates and emergent life history traits respond to fire regimes. We then quantify the extent to which vital rates and life history traits drive demographic resilience. We hypothesize that: (H2) more severe fire regimes decrease populations’ resilience (*i.e.,* lower resistance, lower compensation, and slower recovery); (H3) species’ populations with greater generation time will have higher resistance but slower recovery time; and (H4) populations with greater reproductive output, higher body growth rate, and higher thermal performance will have higher compensation and faster recovery time.

## 2 MATERIALS AND METHODS

### 2.1 Population monitoring

Our study site (*Reserva Ecológica do IBGE*, RECOR; see more details in the SI) is an ideal natural experiment to test our hypotheses regarding the drivers of demographic resilience.

From 1972 to 1990, RECOR was protected from fires. In 1989, a long-term experiment was initiated to evaluate the responses of animals and plants in 10 ha plots (200 x 500 m), each submitted to unique prescribed fire regimes characterized by a combination of timing (early dry season, in late June; mid dry season, in early August; and late dry season, in late September) and frequency (biennial and quadrennial) (Miranda 2010). Plots were placed in the cerrado *sensu stricto* physiognomy and shared a common recent fire history before the experiment’s onset. Thus, differences between plots over time are likely to have arisen due to the prescribed fire regimes (Costa *et al*. 2021).

Here we focus on the three most abundant lizard species from the assemblage to examine the traits that confer higher resilience to weather, microclimate, and fire regimes. Of critical importance to test our hypotheses, each species presents different life history strategies (driven by body sizes, reproductive modes, and clutch sizes). To acquire the individual-level data used in our analyses, from November 2005 to December 2019, we captured individual lizards in study plots using pitfall traps with drift fences in five plots (Fig. S1): three under prescribed biennial fires (early biennial – EB, middle biennial – MB, and late biennial – LB), one under prescribed quadrennial fires (middle quadrennial – Q), and one control plot (C) (Fig. S1). Considering the number of fires each plot was submitted and their canopy cover (Fig. S6), fire severity increased in the following order C < Q < EB < MB < LB. We opened traps for six consecutive days every month and checked them daily during this period. We recorded snout-vent length (SVL) with a ruler (1 mm precision) from each captured individual, followed by marking by toe-clipping (after the appropriate bioethics approval 33786/2016) and release at the capture site. Toe-clipping is a widely used system to mark individuals in amphibians and reptiles, where each toe has a unique code to set the identity of each individual (Hero 1989). We also assessed the reproductive state by palpation in females. This assessment is possible because when female lizards bear eggs, embryos, or enlarged follicles, they display a significant enlargement in their abdomens (Meiri, Brown & Sibly 2012).

Our approach considers that every plot comprises individual populations of each species, where individuals are subjected to different environments promoted by the long-term effects of fire regimes. Immigration and emigration, *i.e.*, movement into and movement out of a study area of arbitrary size and location, *sensu* Fryxell, Sinclair, and Caughley (2014), may bias vital rates estimates and the underlying effects of the experimental plots (Paquet *et al*. 2020), here fire regimes. However, considering the distance between transects in adjacent plots (*ca.* 200 m; Fig. S1A), the small size, and the territorial behavior of some species (Van Sluys 1997), we assumed no significant lizard movements among plots that could affect our results. Indeed, there were no inter-plot recaptures, supporting our assumption. In addition to the costs and difficulties of replication typical of large-scale ecological studies, our experiment could not be replicated because of legal issues associated with burning the vegetation inside protected areas. While this approach may reduce the statistical power of our study, doing so is the only way to ensure the availability of adequate treatment levels in such a large design (Driscoll *et al*. 2010).

### 2.3 Ecophysiological traits, weather, and microclimate

To test (H1) whether ecophysiological traits and weather/microclimate shape how underlying vital rates and emergent life history traits respond to fire regimes, we estimated locomotor thermal performance curves (TPC) (Huey & Stevenson 1979) and hours of activity (*ha*) (Sinervo *et al*. 2010) based on laboratory experiments from at least 10 individuals by each species (see SI for more details). We initially had 14 microclimatic and weather variables. However, after excluding highly correlated variables (using the stepwise procedure *vifcor* from package USDM), we retained: mean air temperature at 200 cm height (*tmed2m*), minimum air temperature at 0 cm height (*tmin0cm*), mean air temperature at 0 cm height (*tmed0cm*), maximum relative air humidity (*RHmax*), solar radiation (*sol*), and accumulated precipitation (*precip*). The *vifcor* algorithm finds a pair of variables highly correlated (cor > 0.8) and excludes the one with a greater variance inflation factor (VIF) repeatedly until no pair of variables highly correlated remains. We used this approach because one of our primary goals is to understand the specific contributions of individual environmental factors to the vital rates (H1) rather than composite axes of environmental variation (such as from a principal component analysis). With the hourly microclimate temperature estimates at 10 cm height, we predicted monthly mean locomotor performance (*perf*) and the hours of activity (*ha*). SI presents the correlations and relationships among the variables (Fig. S7) and the details of the microclimatic and thermal physiological estimates.

### 2.4 Data analyses

We estimated each species’ monthly vital rates (survival, growth, and reproduction) using different datasets integrated into Bayesian hierarchical models and a combination of Cormack-Jolly-Seber (CJS) and Pradel Jolly-Seber (PJS) mark-recapture models of open populations (Cormack 1964; Pradel 1996; Tenan *et al*. 2014). Because the CJS model allows for individuals’ covariates, we related the survival (*σ*) to the individuals’ body size (SVL), while the PJS incorporated the monthly environmental variation (weather, ecophysiological, microclimate, months with fire, and time since last fire – *TSLF*) into populations’ survival (*σ*) and recruitment (probability of recruitment, *p_rec_*). The recruitment parameter (*p_rec_*) is based on the total population (males and females) and can be interpreted as mostly hatchling recruitment since their peaks coincide with captures of small-sized hatchlings (Fig. S8). Additionally, our mark-recapture models are sex-averaged since we have not found differences in survival and growth between males and females (Fig. S9). For the changes in the body size through time (growth, *γ*), we combined a Von Bertalanffy body growth function with the individuals’ records of SVL and capture history. For reproduction, we considered three different processes: the probability of reproduction (*p_rep_*), the production of newborns (clutch size; *n_b_*), and the probability of recruitment (*p_rec_*). We used Bayesian generalized linear models to relate *p_rep_* and *n_b_* to the individuals’ SVL. We implemented the models for each species with JAGS in R, using the packages JAGSUI, RUNJAGS, and RJAGS (Kellner 2019; Plummer 2019). SI presents the details of the parameterizations estimated in each model, including their formulas.

To test H2, H3, and H4 (see Introduction), we used the vital rates (*σ*, *γ*, *p_rep_*, *n_b_*, and *p_rec_*) parameter estimates to build stochastic Integral Projection Models (IPMs) using the IPMR package (Levin *et al*. 2021). Although we have not parameterized the IPMs with explicit density-dependent parameters constraining their vital rates, we assumed that the environmental covariates and plots represented the primary resources the lizards use (habitat, thermal niche, and food), thus indirectly modeling density dependence; this is a common approach when analyzing long-term demographic data (Salguero-Gómez *et al*. 2016a), as it is our case. IPMs are structured population models where the vital rates are influenced by a continuously varying phenotypic trait (*e.g.*, body size) instead of discrete stages or ages (Easterling, Ellner & Dixon 2000). In a population structured by a continuous variable body size (SVL), as in our three study species, the IPM describing the number of individuals of size *y* at time *t* + 1 is:

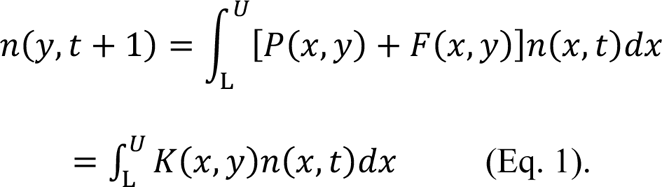

In equation 1, *L* and *U* specify the lower and upper body size limits in the model, respectively; *P*(*x,y*) describes the changes in size conditional on survival; *F*(*x,y*) quantifies fecundity; and *K*(*x,y*) *= P*(*x,y*) *+ F*(*x,y*) is the *kernel* surface, representing all possible transitions and per-capita sexual contributions from size *x* to size *y* between *t* and *t*+1. The *P* sub-kernel incorporates both survival and growth components, such that *P*(*x,y*) *= σ*(*x*) × *γ*(*x,y*), where *σ*(*x*) is the survival probability of a size-*x* individual and *γ*(*x, y*) is the probability of a size-*x* individual growing to size *y* (see SI for the IPMs’ implementation). Considering the five plots and 169 months, excluding the last month that mark-recapture models cannot estimate (Cormack 1964; Pradel 1996), we generated 845 monthly IPM kernels for each species.

To test whether populations’ generation time (H3) and reproductive output (H4) shape their demographic resilience, we estimated those two life history traits with the monthly IPM kernels of each plot and species using the package RAGE (Jones *et al*. 2022). We also estimated other life-history traits from these IPMs, but we decided to focus our analyses on generation time and net reproductive output only, as they describe the principal axes of the life history variation in our species (Figs. S13-S14), as well as in other animals (Gaillard *et al*. 2005; Healy *et al*. 2019), including reptiles (Rodriguez-Caro *et al*. 2023).

To test our hypotheses regarding the demographic resilience components (H2-H4), we estimated the resistance, compensation, and recovery time from our IPMs. To do so, we used the package POPDEMO (Stott, Hodgson & Townley 2012) on our monthly, species- and site-specific IPMs. We used the first time-step amplification (reactivity) and attenuation as the measures for compensation and resistance, respectively, following Capdevila *et al*. (2020). First time-step amplification and attenuation are the highest population increase and the lowest population decrease possible in the next month, respectively. Attenuation values vary from 0 to 1, where 0 means low resistance and 1 means high resistance. Thus, given the species, sites, and monthly environmental covariates (weather, microclimate, ecophysiological, fire-related, and randomness), resistance and compensation estimates represent the monthly potential of each population to maintain and increase its size in the next month, respectively. Recovery time was estimated as follows:

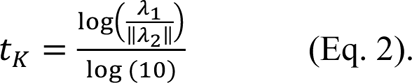

In equation 2, λ_1_ and λ_2._ correspond to the dominant and subdominant eigenvalues of the *K* kernel, respectively. This measure estimates the time (months) required to reach a stable stationary structure every month.

Next, to test (H2) whether more severe fire regimes decrease populations’ resilience populations, and populations with (H3) greater generation time have higher resistance but slower recovery, and (H4) populations with greater reproductive output have higher compensation and faster recovery, we built Bayesian dynamic generalized additive models using MVGAM (Clark & Wells 2023). We related the demographic resilience components with the two life history traits and among plots. We used gamma error distributions for compensation and recovery time, and for resistance, we used a beta distribution. For each population time series (by plot and species), we included plot as a random variable to control for pseudoreplication and non-independency among plots and repeated observations. We also included a latent autoregressive and moving-average trend model (*AR1MA*) to capture the first-lag autocorrelation. We log_10_-transformed the compensation values and z-scaled the life history traits values prior to analyses.

To test specific aspects of H4, that populations with higher body growth rate and higher thermal performance will have higher compensation and faster recovery time and, at the same time, inspect which IPM vital rate parameters affected the different demographic resilience components, we conducted a brute-force sensitivity analysis (Morris & Doak 2002). To implement the brute force sensitivity analysis, we added values from 0.01 to 0.1 to each of the parameters specifying our vital rates and recalculated the differences in the demographic resilience components. Our parameter sensitivities (*S_p_*) were estimated as in equation 3, where the numerator describes the difference in the transient metric of choice (*x*; *e.g.*, reactivity for amplification) resulting from perturbing a vital rate parameter *p* at a time, in the denominator, following Cant *et al*. (2023):

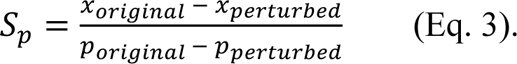

## 3 RESULTS

### 3.1 Vital rates

Overall, we found that weather and microclimate shape the survival and recruitment of *Micrablepharus atticolus* and *Tropidurus itambere*, but not of *Copeoglossum nigropunctatum* (Fig. 1), partially supporting our hypothesis (H1). As opposed to H1, ecophysiological traits were poor predictors of the vital rates, except for the recruitment of *M. atticolus*. For instance, the vital rates of survival and recruitment of *C. nigropunctatum* were better explained by random monthly variation than environmental covariates (Fig. 1; Table S4). In contrast, survival of *M. atticolus* and *T. itambere* was only affected by weather (*precip*, *tmed2m*, and *sol*; Fig. 1; Table S4). Partially supporting our predictions (H1), weather (*tmed2m*), microclimatic (*tmin0cm*), and ecophysiological (*perf* and *ha*) conditions affected the recruitment of *M. atticolus* (Fig. 1; Table S4); Recruitment of *T. itambere* only varied with weather conditions (*tmed2m* and *sol*; Fig. 1; Table S4). These patterns resulted in higher variation in the life-history space in these two species—*M. atticolus* and *T. itambere* (Fig. S13). We found differences in these vital rates among plots, which had been exposed to prescribed fire regimes for 18 years (Table S4). The differences in vital rates were most apparent under both extremes of fire severity (absence of fire and frequent late-dry season fires), suggestive of the long-term effects of the fire regimes (Table S4), in addition to differences in microclimate and ecophysiological traits (H1). However, the other fire-related variables (*fire* and *TSLF*) were poor predictors of the vital rates, except for the recruitment of *M. atticolus* (Fig. 1; Table S4). See SI for mark-recapture summary statistics.

**Fig. 1.**
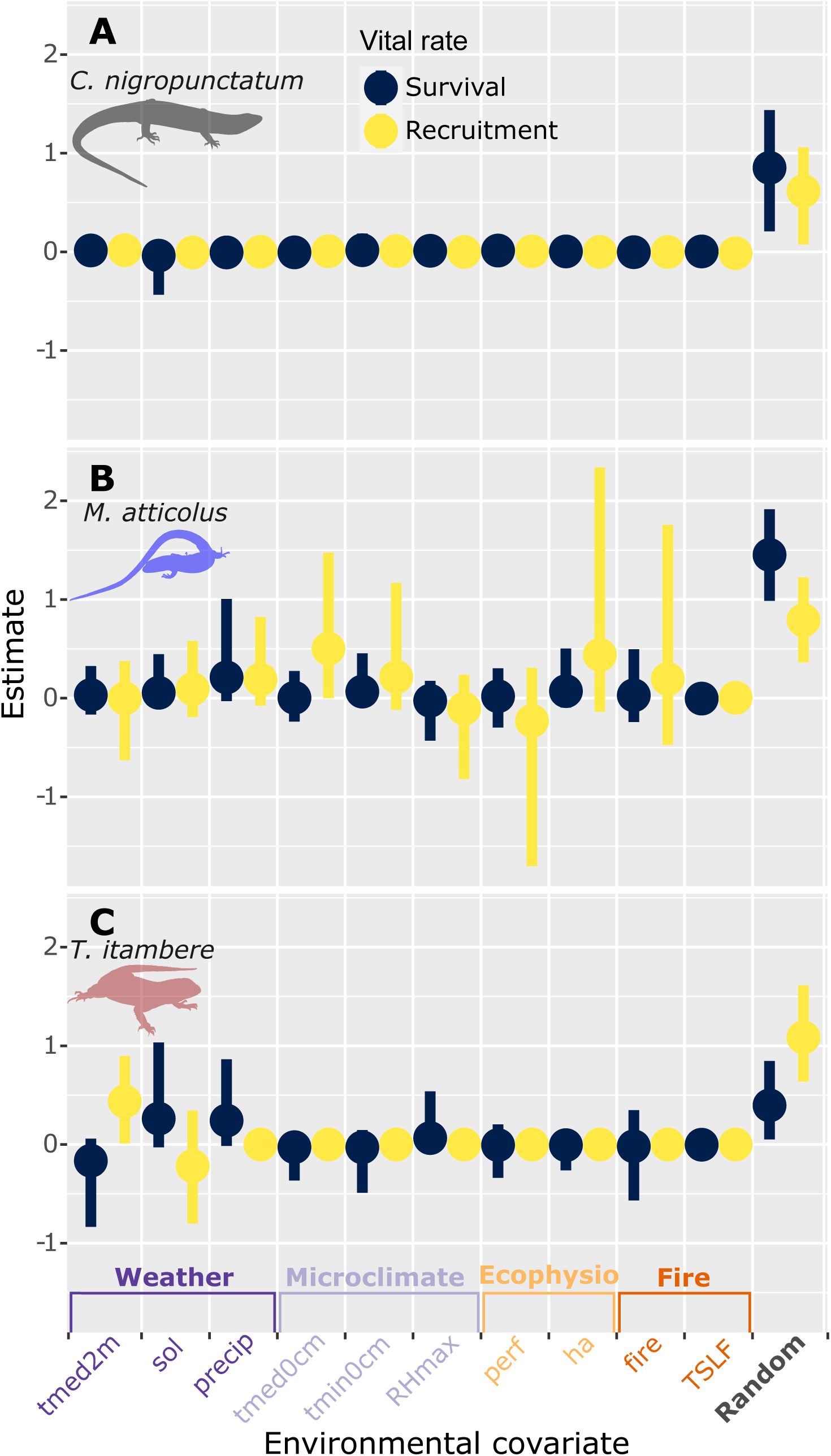
Model coefficients (mean and 95% credible intervals) of the effects of environmental covariates on vital rates of three species of lizards corresponding to *tmed2m* = mean air temperature at 200 cm height; *tmin0cm* = minimum air temperature at 0 cm height; *tmed0cm* = mean air temperature at 0 cm height; *RHmax* = maximum relative air humidity; *sol* = solar radiation; *precip* = accumulated precipitation; *perf* = mean locomotor performance; *ha* = hours of activity; *fire* = month with fire; *TSLF* = time since last fire; *random* = monthly random variation. We group-colored the covariates related to weather, microclimate, ecophysiological traits, fire, and randomness. Species’ silhouettes from photos taken by Nicolás Pelegrin (*C. nigropunctatum* and *M. atticolus*) and Carlos Morais (*T. itambere*).

### 3.2 Fire regimes and demographic resilience

Contrary to H2, populations of the three species presented high resistance ability in severe fire regimes (Figs. 2-3). However, populations of *T. itambere* displayed low compensation in the most severe fire regime (Figs. 2-3). On average, the three lizard species recover rather fast to their stable stationary structure (Figs. 2-3; *C. nigropunctatum* - 1.844 ± 0.064 S.D. months; *M. atticolus* - 1.516 ± 0.291 S.D. months; *T. itambere* - 1.946 ± 0.200 S.D. months). However, they followed clearly differentiated strategies depending on the fire severity, resulting in the lowest resistance, highest compensation, and fastest recovery abilities at intermediate levels of severity (third and fourth levels–frequent early- and mid-dry season fires; Fig. 3). The most striking differences, though, occurred at the extreme of fire severity (frequent late-dry season fires): *M.atticolus* displayed higher resistance and compensation, and *T. itambere* higher compensation and lower resistance (Fig. 3). Interestingly, *M. atticolus* and *T. itambere* also lost the ability to recover fast (*i.e.*, < 1.5 months) from fire and climate disturbances in the most severe fire regime (Fig. 3).

**Fig. 2.**
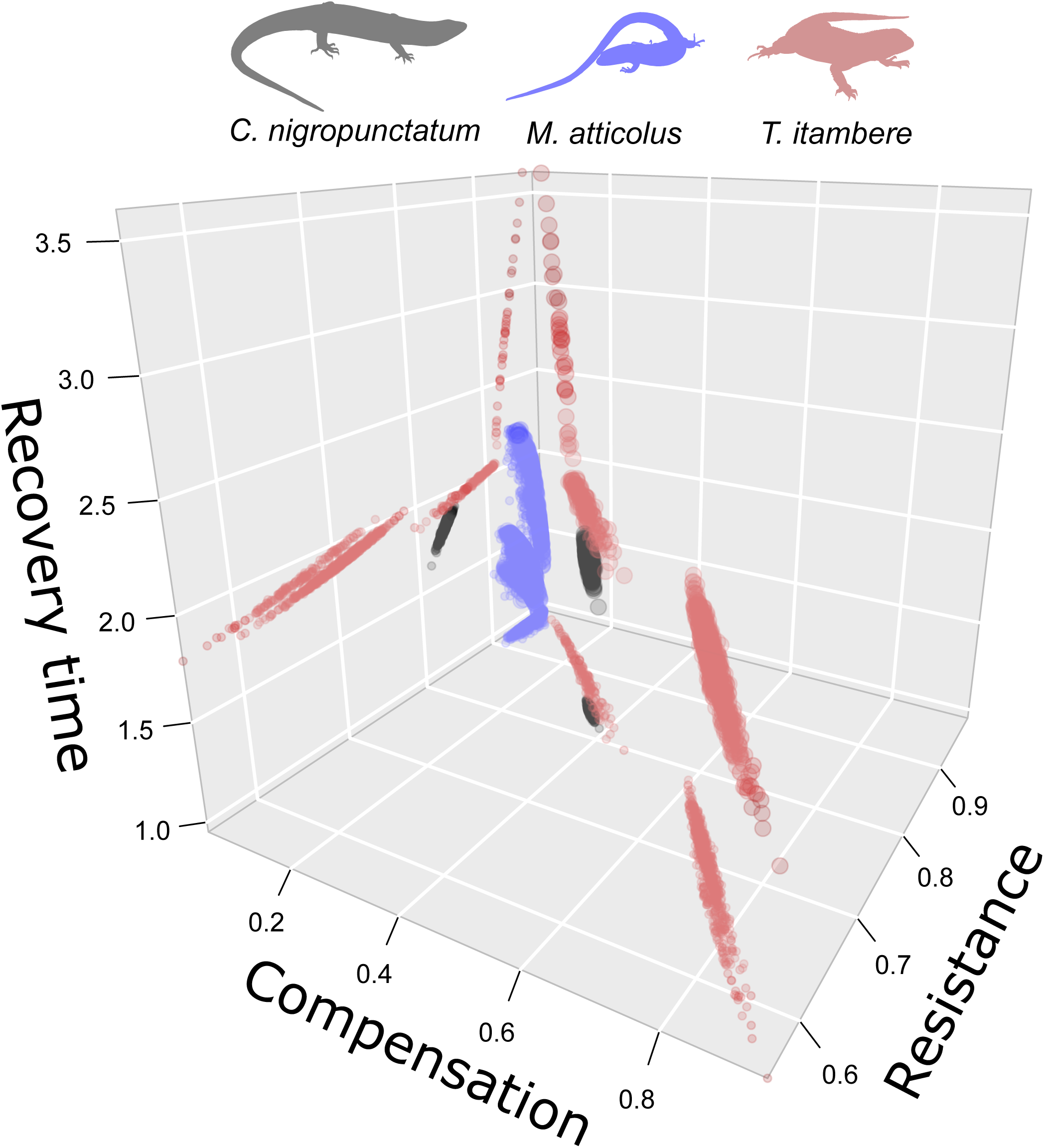
Three-dimensional space of the demographic resilience components (resistance, compensation, and recovery time) for three species of lizards from the Brazilian Cerrado savannas, showing distinct demographic resilience strategies. With a 14-year mark-recapture dataset, we derived demographic resilience components from monthly stochastic Integral Projection Models shaped by weather, microclimate, ecophysiological traits, and fire covariates.

**Fig. 3.**
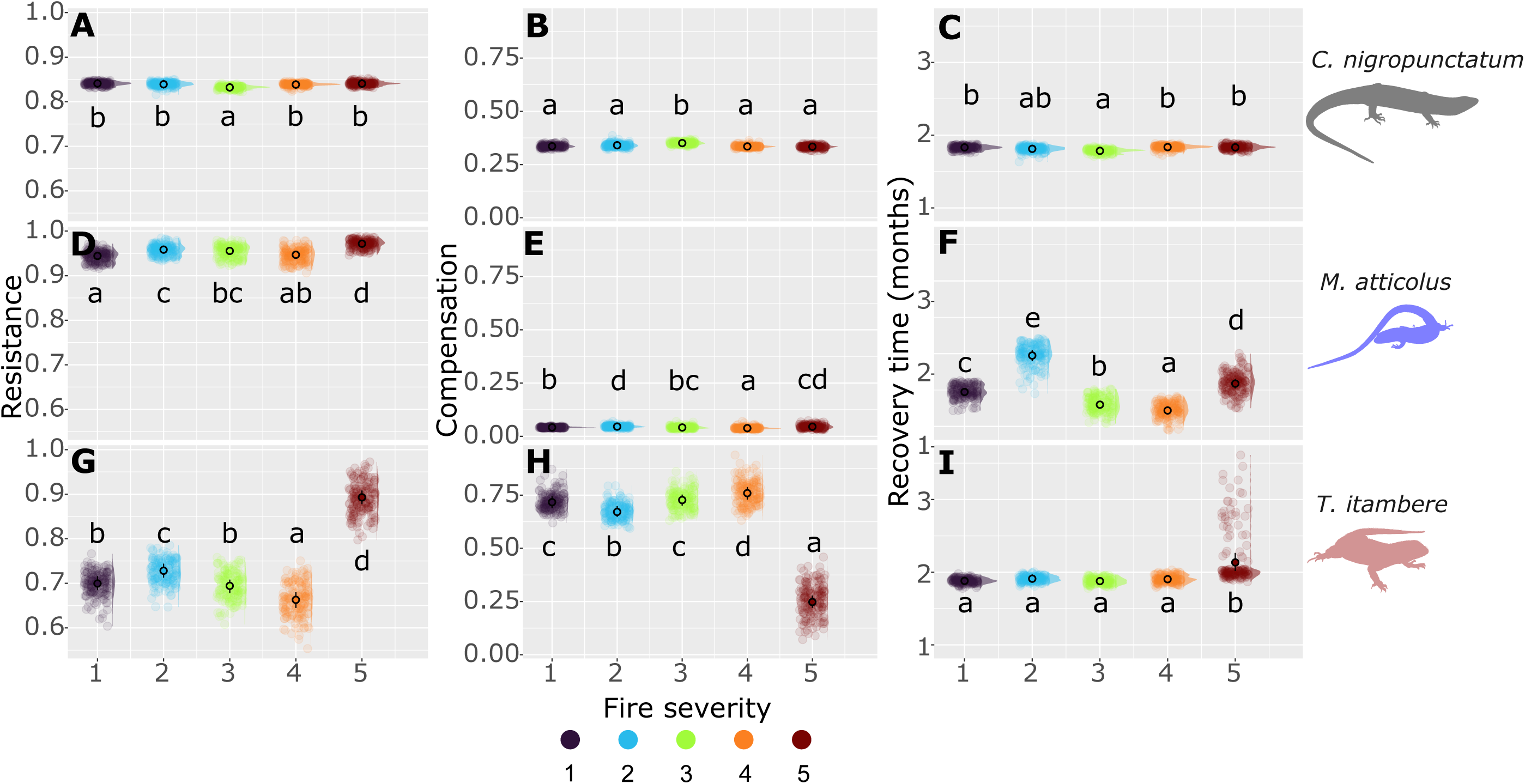
Raw data, density, mean estimates, and 95% credible intervals of the demographic resilience components: (A, D & G) resistance; (B, E & H) compensation, and (C, F & I) recovery time of three species of lizards: (A-C) *Copeoglossum nigropunctatum*, (D-F) *Micrablepharus atticolus*, and (G-I) *Tropidurus itambere* from the Brazilian Cerrado savannas under different fire regime severity. Within each panel, we show high posterior density (95% probability) contrast letters corresponding to groups that are not significantly different (overlapping zero difference). Later letters in the alphabet mean higher values (e > d > c > b > a).

The fire regime effects on the demographic resilience components highly agree with the realized population growth rates and abundance trends in the three species. All populations had mean positive population growth (*log(*λ) > 0) throughout the 14 years of study (Fig. S11). However, *C. nigropunctatum* displayed lower stability (higher S.D. in *log(*λ)) in the intermediate level of fire severity (third level–frequent early-dry season fires), while *M. atticolus* and *T. itambere* populations varied more (higher S.D. in *log(*λ)) in the most severe fire regimes (fourth and fifth levels, SI; Fig. S11). Overall, *C. nigropunctatum* had fewer captures in intermediate fire severity levels (third and fourth levels–frequent early- and mid-dry season fires), while *M. atticolus* and *T. itambere* displayed the opposite pattern (Fig. S2). Both *M. atticolus* and *T. itambere* had low captures at the extremes of fire severity (absence of fires and frequent late-dry season fires; Fig. S2).

### 3.3 Life history traits and demographic resilience

Using Bayesian dynamic generalized additive models, we found that the differences in the demographic resilience components between fire regimes were predicted by generation time and reproductive output (Fig. S15), supporting H3 and H4. However, the direction of some of our predictions differed for the species with fixed clutch size, *M. atticolus*. We found that, in populations with higher generation times, resistance was higher, compensation was lower, and recovery was longer (Fig. 4). This striking pattern was consistent across the three examined species but with a weaker relationship in *C. nigropunctatum* due to its constrained vital rate variation in response to the environment, and thus lower variance in life history traits (see Figs. S13-S15 for different life history traits among the species). An important exception to the overall pattern was again *M. atticolus*, with its fixed clutch size, where populations with higher generation times displayed lower resistance to fire and climate variation.

**Fig. 4.**
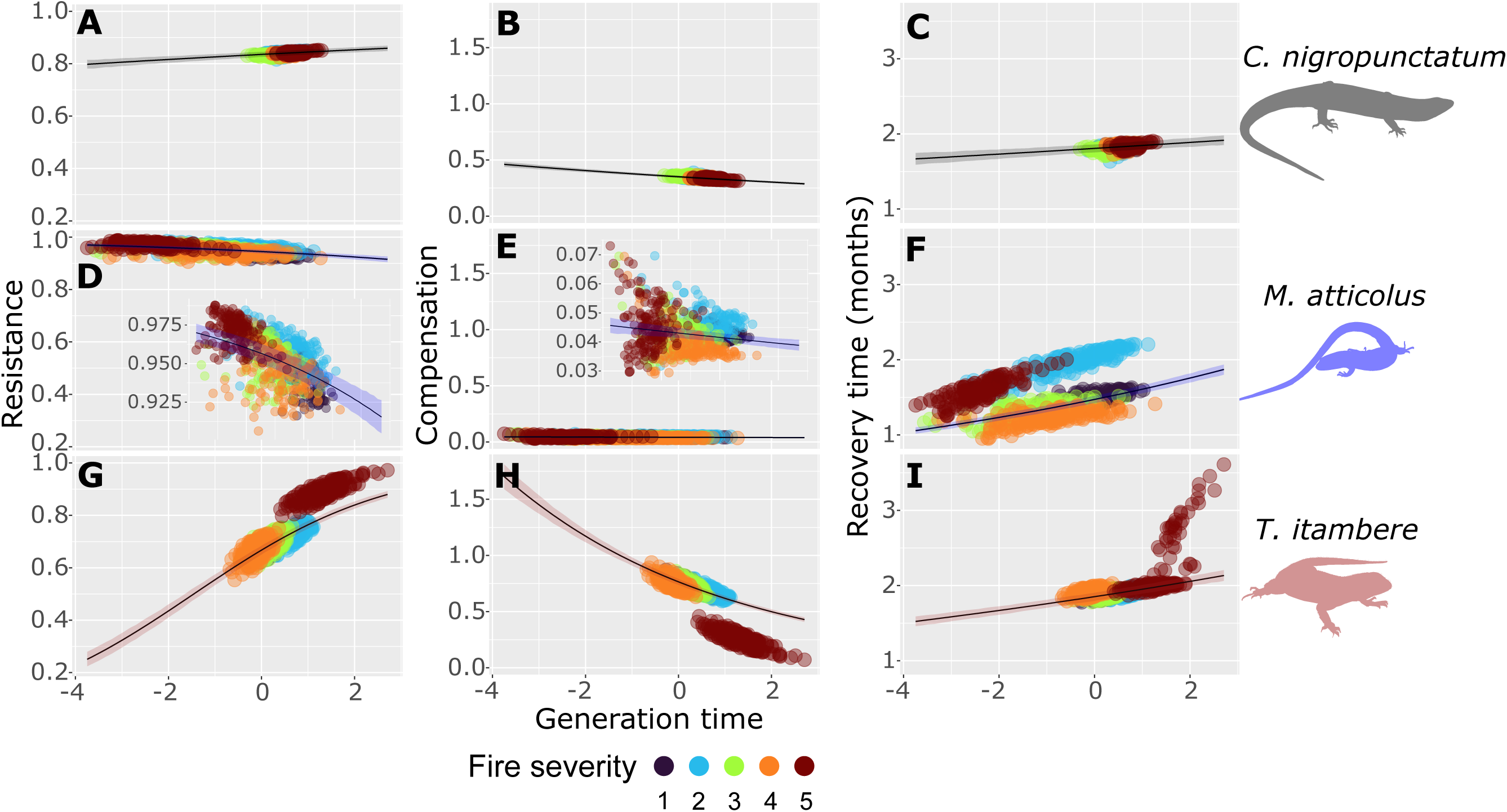
Relationship between generation time (z-scaled) and the demographic resilience components: (A, D & G) resistance, (B, E & H) compensation, and (C, F & I) recovery time of three lizard species: (A-C) *Copeoglossum nigropunctatum*, (D-F) *Micrablepharus atticolus*, and (G-I) *Tropidurus itambere* under fire regimes of varying severity. Lines and shades represent mean estimates and 95% credible intervals derived from Bayesian dynamic generalized additive models, respectively. To offer a higher resolution for *M. atticolus*’ resistance and compensation, we also provide inserts in D and E with a narrower y-axis range.

Reproductive output also shaped the demographic resilience components of the three species, supporting H4. The opposite pattern regarding the relationship between resilience components and life history traits was found with reproductive output (Fig. 5). Indeed, in populations with higher reproductive output, resistance was lower, compensation was higher, and recovery was faster (Fig. 5). This pattern was consistent across species, with the exception of the fixed-clutch species *M. atticolus*, displaying higher resistance and lower compensation with increasing reproductive output (Fig. 5).

**Fig. 5.**
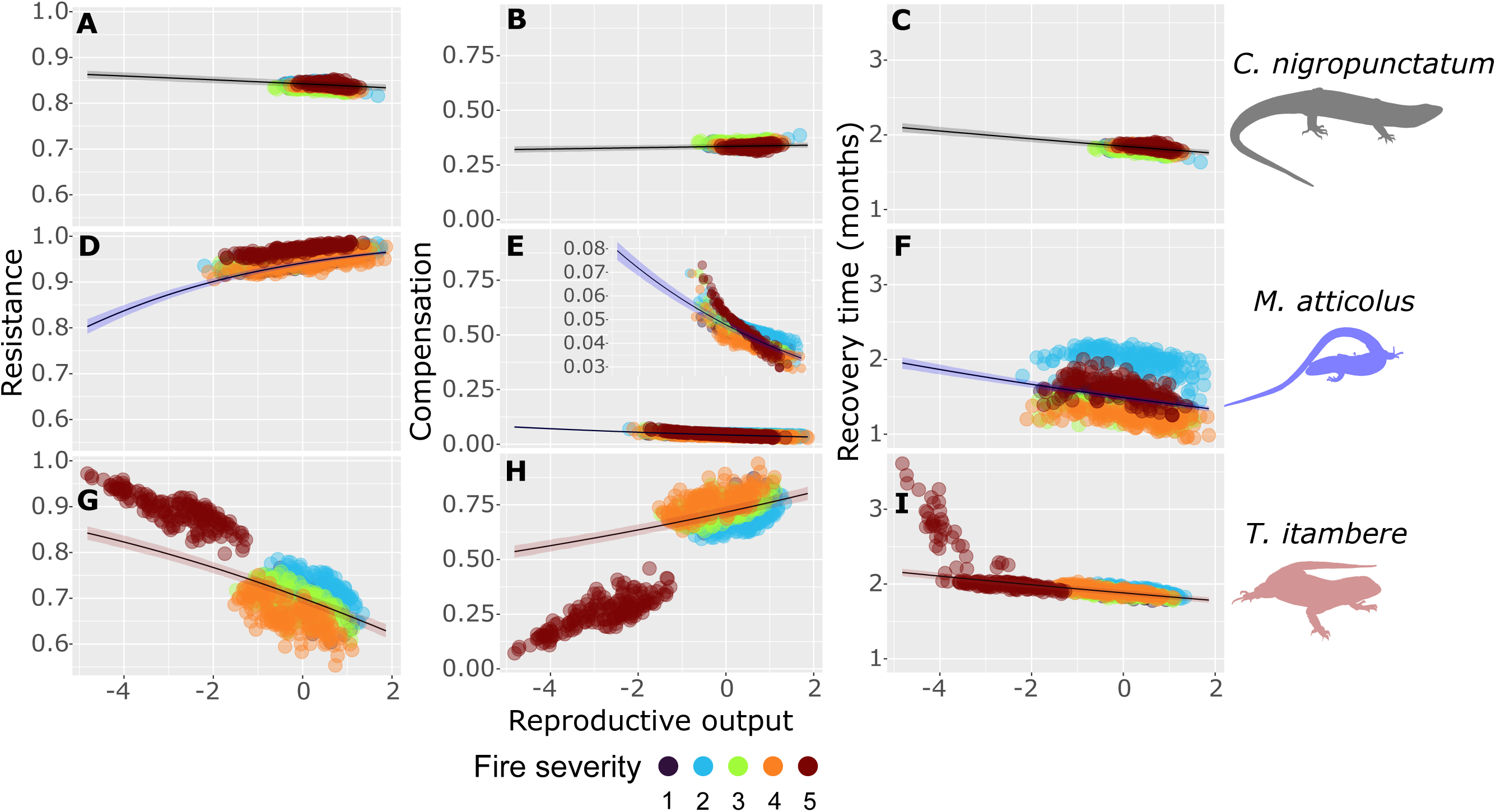
Relationship between reproductive output (z-scaled) and the demographic resilience components (A, D & G) resistance, (B, E & H) compensation, and (C, F & I) recovery time of three lizard species (A-C) *Copeoglossum nigropunctatum*, (D-F) *Micrablepharus atticolus*, and (G-I) *Tropidurus itambere* under fire regimes of varying severity. Lines and shades represent mean estimates and 95% credible intervals derived from Bayesian dynamic generalized additive models, respectively. To offer a higher resolution for *M. atticolus*’ compensation, we also provide an insert in E with a narrower y-axis range.

### 3.4 Vital rate parameters and demographic resilience

The brute-force sensitivity analysis revealed that the demographic resilience components are most affected by the reproductive parameters (Table 1) and, to a lesser extent, by body growth; but not by thermal physiology as we predicted (H4). The relationships between the probability of reproduction or the number of newborns and SVL regulated all the resilience components in the three species (Table 1). This result indicates that reproduction among the largest individuals (usually the oldest, (Halliday & Verrell 1988)) is critical for the demographic resilience of *C. nigropunctatum* and *T. itambere*. Higher vital rate parameter values describing allocation in reproduction led to lower resistance and higher compensation (Table 1). However, complex results emerged regarding the vital rate parameters shaping recovery time. In *T. itambere*, a higher probability of reproduction and a higher number of newborns resulted in a displacement of the population structure further away from the stable size distribution, leading to longer recovery times (Table 1). In *C. nigropunctatum*, we found the opposite pattern, except for the probability of reproduction (which presented complex non-linear relationships). In *M. atticolus,* a higher probability of reproduction of larger adult individuals shortened recovery times (Table 1). As we predicted, slower body growth rates led to slower recovery time, especially for the small-sized *M. atticolus* (Table 1). Other parameters had negligible effects on the demographic resilience components across species and fire regimes.

**Table 1.** Sensitivity (mean ± S.D.) of the demographic resilience components to parameters’ perturbation in three species of lizards from the Brazilian Cerrado savannas: *Copeoglossum nigropunctatum*, *Micrablepharus atticolus*, and *Tropidurus itambere*. Cell colors vary from white to dark orange, indicating absolute sensitivity values of demographic resilience component concerning different plausible drivers: α_vitalrate_ = intercept of the vital rate function; α_vitalrateSVL_: slope of the linear relationship with body size (snout-vent length— SVL); β_vitalrateSVL_^2^: slope of the quadratic relationship with body size (SVL); μ_K_ = mean body growth constant, *i.e.*, time required to reach the asymptotic size, thus higher μ_K_ values means slower body growth rates. Ony parameters related to probability of reproduction (*prep*), number of newborns (*nb*), and body growth were higher than zero. The sensitivity values of the other vital rates were null and not shown. Negative values, which mean negative effects on the demographic resilience components, are biologically meaningful and mathematically possible.

| Parameter | Resistance | Compensation | Recovery time |
| --- | --- | --- | --- |
| *Copeoglossum nigropunctatum* |  |  |  |
| $\alpha_{\text{prep}}$ | -0.026 ± 0.002 | 0.111 ± 0.004 | 0.052 ± 0.001 |
| $\beta_{\text{prepSVL}}$ | -2.909 ± 0.315 | 13.836 ± 0.784 | 8.023 ± 0.264 |
| $\beta_{\text{prepSVL}}^2$ | -71.366 ± 69.583 | 86.673 ± 102.262 | -173.243 ± 108.166 |
| $\alpha_{\text{nb}}$ | -0.067 ± 0.002 | 0.2 ± 0.001 | -0.21 ± 0.012 |
| $\beta_{\text{nbSVL}}$ | -9.58 ± 0.219 | 27.562 ± 1.194 | -6.326 ± 0.745 |
| $\mu_K$ | 0.007 ± 0.001 | 0.004 ± 0.001 | 0.03 ± 0.005 |
| *Micrablepharus atticolus* |  |  |  |
| $\alpha_{\text{prep}}$ | -0.037 ± 0.011 | 0.035 ± 0.005 | -0.103 ± 0.182 |
| $\beta_{\text{prepSVL}}$ | -1.475 ± 0.462 | 1.603 ± 0.219 | -4.227 ± 8.933 |
| $\mu_K$ | $0.012 \pm 0.003$ | $0.005 \pm 0.001$ | $0.654 \pm 0.251$ |
| *Tropidurus itambere* |  |  |  |
| $\alpha_{\text{prep}}$ | $-0.036 \pm 0.008$ | $0.127 \pm 0.02$ | $0.059 \pm 0.005$ |
| $\beta_{\text{prepSVL}}$ | $-2.740 \pm 0.608$ | $10.516 \pm 1.557$ | $5.573 \pm 0.342$ |
| $\alpha_{\text{nb}}$ | $-0.128 \pm 0.005$ | $0.296 \pm 0.003$ | $-0.033 \pm 0.004$ |
| $\beta_{\text{nbSVL}}$ | $-8.806 \pm 0.201$ | $24.955 \pm 0.319$ | $5.312 \pm 0.914$ |
| $\beta_{\text{nbSVL}}^2$ | $-186.827 \pm 141.564$ | $1655.239 \pm 163.511$ | $3.747 \times 10^{10} \pm$<br>$9.308 \times 10^{10}$ |
| $\mu_K$ | $0.008 \pm 0.006$ | $0.005 \pm 0.004$ | $0.005 \pm 0.004$ |

## 4 DISCUSSION

Using 14 years of monthly mark-recapture data of three lizard species from the Brazilian Cerrado savannas, we show that severe disturbance regimes result in overall less resilient populations, supporting our primary hypothesis (H2). Indeed, our stochastic IPMs demonstrate that high-severity fire regimes increase resistance but decrease compensatory or recovery abilities, leading to less stable populations. Accordingly, we also show that generation time and reproductive output predict the differences in the demographic resilience components (resistance, compensation, recovery time) among fire regimes and species. Here, populations with higher generation times are characterized by a greater degree of resistance and slower recovery time (H3), whereas those with higher reproductive output tend to compensate more and recover faster (H4). The only species with different relationships between these life history traits and demographic resilience components is the small body-sized lizard *Micrablepharus atticolus,* which has a fixed clutch size of two eggs. Despite these differences, all species present a strong negative correlation between resistance and compensation to fires and climatic stochasticity. Partially supporting our hypothesis, the species’ probability and quantity of reproduction, but not its ecophysiology, are the main drivers of the demographic resilience in our three ectothermic species (H1), with body growth only being a strong predictor of recovery time in *M. atticolus*.

The most severe fire regime results in more resistant populations in our three species. However, our lizards display longer recovery times (∼ two months) or lower compensation in response to this severe fire regime. This finding suggests that these populations surpass a tipping point in severe fire regimes and achieve an alternative stable state to persist, as shown in other systems, such as in shrubs and grasses from tallgrass prairies (Collins *et al*. 2021). Here, the new state after intense fires is characterized by strategies that allocate resources to either side along the fast-slow continuum (Stearns 1976) or the reproductive strategies continuum (Salguero-Gómez *et al*. 2016b). For instance, populations of *T. itambere* display longer generation times but lower reproductive output in the most severe fire regime. Similarly, *M. atticolus* also presents lower reproductive output but a faster pace of life (*i.e.*, lower generation time) in this regime. These findings further highlight the plasticity of species resilience strategies to cope with different levels of disturbance so populations may remain viable, an area that has received considerable attention in recent years (Vinton *et al*. 2022). Said plasticity seems particularly high in reptiles, which have not diminished their ability to respond to disturbances in the last decades compared to other vertebrates, such as mammals and birds (Capdevila *et al*. 2021a).

Our findings align with recent examinations of the relationship between the demographic resilience of plants and animals with the fast-slow continuum (Stott *et al*. 2010; Jiang *et al*. 2022). Indeed, recent studies have recently shown that populations with longer generation times tend to display a lower ability to compensate and are slower to recover after disturbances (Stott *et al*. 2010). In addition, here we show a clear relationship between demographic resilience and reproductive strategy, an orthogonal axis to the fast-slow continuum (Salguero-Gómez *et al*. 2016b; Healy *et al*. 2019). For instance, our populations with higher reproductive output resist less, compensate more (except the species with fixed clutch size, *M. atticolus*), and recover faster after disturbances. This reactive response to disturbances agrees with work showing that the ability to compensate, linked to reproductive output too, enhances colonizing and invasive potential among reptiles and amphibians (Allen, Street & Capellini 2017), and even in plants (Iles *et al*. 2015).

Multiple calls have been made for a more mechanistic understanding of resilience in ecological systems (Hodgson, McDonald & Hosken 2015; Kefi *et al*. 2019; Capdevila *et al*. 2020); these calls remain largely unanswered (Capdevila *et al*. 2021b). Here, we contribute to this knowledge gap by mechanistically integrating into the local population dynamics of each species their ecophysiology, size structure, environmental factors, and varying disturbance regimes. Indeed, one of our key findings is that probability and quantity of monthly reproduction are key drivers of demographic resilience. The lack of predictive power of ecophysiological characteristics (*e.g.*, thermal performance, preferred temperature) on resilience reinforces the idea that life history traits constitute an excellent approach to understanding demographic resilience, as recently highlighted in a synthesis across 69 animal and 232 plant species (Capdevila *et al*. 2022). In addition, our results suggest that resilience components exhibit strong intra- and inter-specific variation and that these relationships are strongly modulated by species’ vital rates and life history characteristics. Thus, we argue that existing macroecological and comparative approaches (Capdevila *et al*. 2022) likely mask some of the intraspecific trade-offs we disentangle here using 14 years of monthly mark-recapture data under five levels of fire disturbances. Future studies will likely deliver important insights by testing the predictive power of various kinds of traits (*e.g*., morphological, physiological, life history) across a wider range of species and disturbance levels.

Fast growth increases resilience after disturbances (Pausas & Bond 2020) and enhances persistence in novel environments (Lytle 2001; Allen, Street & Capellini 2017). In line with these findings, we show that strategies leading to earlier reproduction are effective at conferring compensation and faster recovery against fires and climate variation in our three lizard species. We suggest that this strategy may be particularly important for small animals, as it may allow them to achieve sexual maturity earlier, like in the case of *M. atticolus*. In contrast, we show that in months when individuals allocate more resources to reproduction, populations lose the ability to resist disturbances. This pattern presumably results from individuals’ reproductive costs (Schwarzkopf & Shine 1992); thus, by delaying reproduction, individuals of indeterminate growth—most reptiles (Halliday & Verrell 1988)—can continue to increase survival and reproductive rates at older ages (Halliday & Verrell 1988; Healy *et al*. 2019), buffering the population against disturbances and environmental stochasticity (Koons, Metcalf & Tuljapurkar 2008). The timing of disturbances may affect population resilience and explain why the timing of reproduction (*i.e.*, phenology) has a critical effect on the responses of lizards (Braithwaite 1987) and plants (Zirondi, Ooi & Fidelis 2021) to fires prescribed in different seasons. This connection between phenology and the timing of disturbances is increasingly relevant to other highly seasonal environments, where species’ vital rates are adapted to predictable environmental changes (Prather *et al*. 2023).

Since reproduction is a critical vital rate conferring resilience, species with morphological and physiological constraints on reproduction (*e.g.*, viviparity and fixed clutch sizes) have limited ability to benefit from disturbances. This prediction is precisely what we show in *C. nigropunctatum* and *M. atticolus*. However, both species relatively sustained their resilience strategies in varying degrees of fire severity. Recently reported links between demographic buffering and transient dynamics after disturbances (demographic resilience) (Capdevila *et al*. 2020) may explain these patterns (Stott *et al*. 2010; McDonald *et al*. 2016). In reptiles, viviparity developed as an adaptation to increase reproduction success in cold climates (Zimin *et al*. 2022), and might also confer species with the ability to buffer populations against environmental stochasticity, including resilience to disturbances. Therefore, demographically buffering species may outcompete others in the most extreme fire regimes (total absence of fire *vs.* frequent and severe fires), as we show here. However, because of the constraints on their reproductive schedules, individuals from viviparous and fixed clutch-size species need to perform other strategies to increase their reproductive output and to compensate after fires, such as reproducing more often (Cayuela *et al*. 2022). Indeed, these results provide the possible mechanisms for why, among 75,000 species of vascular plants and tetrapods, long-lived and low-reproductive species present higher extinction risks than fast-living and high-reproductive ones (Carmona *et al*. 2021).

Our findings have key management and conservation implications. First, easily measurable traits such as probability of reproduction and clutch size predict species’ resilience to environmental stochasticity and fires. Despite being more resistant, species with high generation time and low reproductive output take longer to recover and cannot compensate as much as species with faster paces of life. In our study, populations have higher resilience at low and intermediate degrees of fire severity. Therefore, these results support the pyrodiversity hypothesis, whereby higher heterogeneity in fire regimes increases the resilience of different populations and communities and avoids high-severity regimes that homogenize the environment (Jones & Tingley 2021). Second, despite species persisting in highly severe fire regimes, populations can attain alternative stable states (Collins *et al*. 2021), with limited compensatory and recovery abilities after disturbances. Thus, traditional ecological community studies probably underestimate disturbance impacts since populations may persist in severe regimes but not in optimal conditions. Third, reproduction is a key mechanism conferring higher compensation and faster recovery time, both in our species and others (Iles *et al*. 2015; Allen, Street & Capellini 2017). Accordingly, species with constraints on reproduction (*e.g.*, viviparity, fixed clutch size) have higher resistance but lower compensation and take longer to recover from disturbances. Taken together, these key findings imply that, to enhance viable strategies after disturbances, managers should focus on generating suitable and heterogeneous habitats to facilitate the reproductive aspects of populations. The ability to quantify and predict the resilience components of natural populations based on their vital rates and life history traits will be increasingly critical for conservation biology in the coming decades when climate change and disturbances are projected to increase in frequency and intensity (Bowman *et al*. 2020).

## Acknowledgments

This study was financed in part by the Coordenação de Aperfeiçoamento de Pessoal de Nível Superior - Brasil (CAPES) - Finance Code 001 and by a NERC Independent Research Fellowship to RSG (NE/M018458/1). HCS was supported by Coordenação de Aperfeiçoamento de Pessoal de Nível Superior – Brasil (CAPES) [grant numbers 88887.484511/2020-00 and 88881.623332/2021-01]. GRC thanks Coordenação de Aperfeiçoamento de Pessoal de Nível Superior (CAPES), Conselho Nacional de Desenvolvimento Científico e Tecnológico (CNPq), Fundação de Apoio à Pesquisa do Distrito Federal (FAPDF), and the USAID’s PEER program under cooperative agreement AID-OAA-A-11-00012 for financial support.

## Statement of authorship

Conceptualization: HCS, AM, GRC, and RSG; Data curation: HCS and GRC; Formal analysis: HCS with assistance by RSG; Funding acquisition: GRC and RSG; Investigation: HCS and GRC; Methodology: HCS, GRC, and RSG; Project administration: GRC; Resources: GRC and RSG; Software: HCS and GRC; Supervision: AM, GRC, and RSG; Validation: AM, GRC, and RSG; Visualization: HCS with assistance by RSG; Writing - original draft: HCS; Writing - reviewing and editing: AM, GRC, and RSG.

## Data availability statement

We confirm that the data and code supporting the results will be archived in an appropriate public repository such as Dryad or Figshare, and the data DOI will be included at the end of the article.

