## Supplementary material for "Severe fire regimes decrease resilience of ectothermic populations": SI

### Supporting Information - Severe fire regimes decrease resilience of ectothermic populations

#### Study area

We conducted this study at Reserva Ecológica do IBGE, RECOR (15°56’41” S, 47°53’07” W; 1,141 m above sea level; Fig. S1), situated approximately 15 km south of Brasília, Distrito Federal, Brazil, in the core of the Cerrado biome. The climate there corresponds to the Aw (tropical with dry winter) type in Köppen’s classification and has a high intra-annual predictability (Alvares *et al.* 2013). Mean temperatures are mild year-round (20.6 ± 1.4 °C S.D.), varying between 18.0–22.4 °C. Conversely, precipitation is markedly seasonal, with a wet season between October and April, when ~94% of the 1417.0 mm mean annual occurs. The vegetation consists of a complex mosaic of savannas, grasslands, and gallery forests (Eiten 1972).

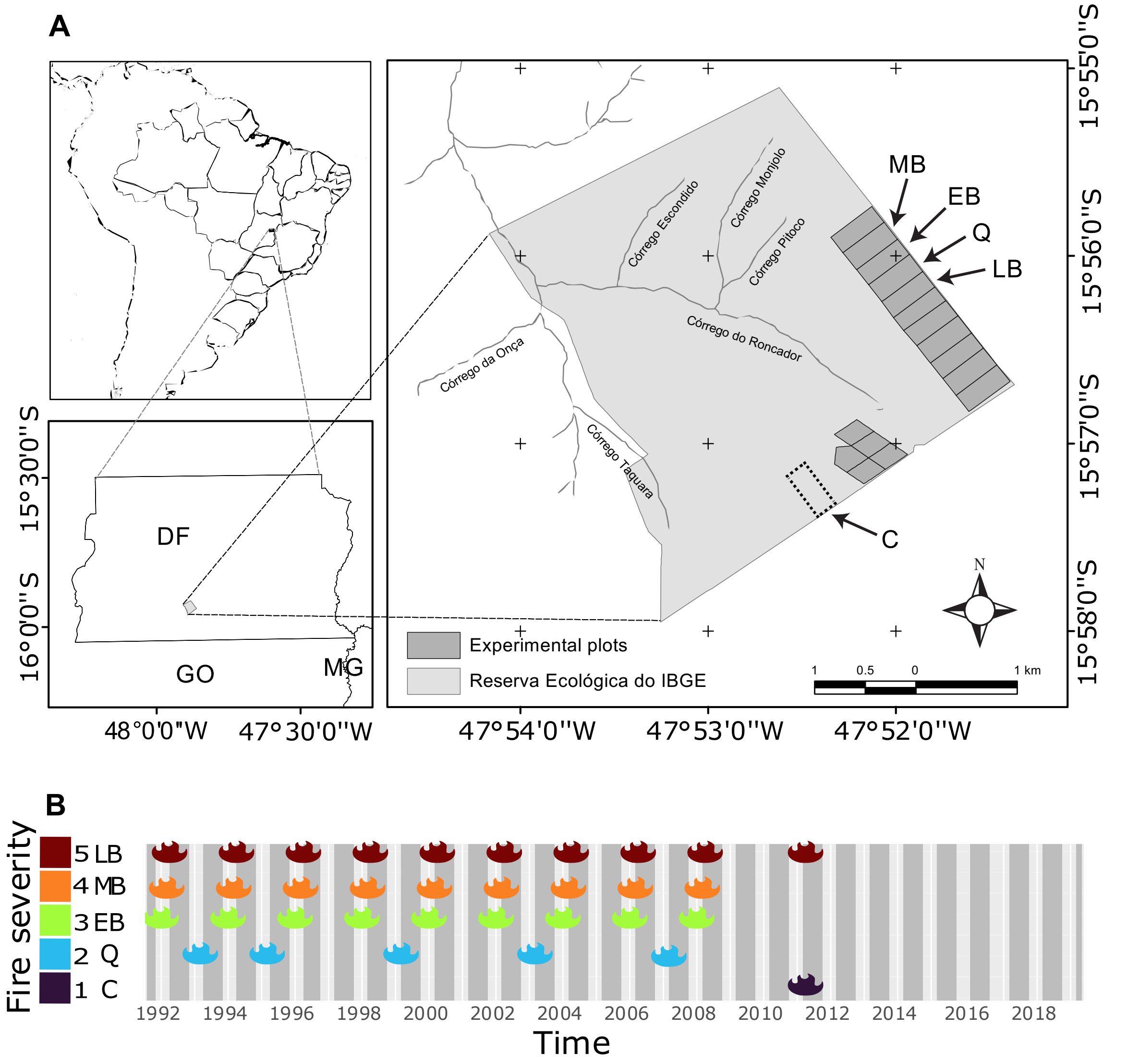


**Fig. S1.** (A) Map with location of the studied plots, each exerted to different fire timings and frequencies: late biennial (LB), middle biennial (MB), early biennial (EB), quadrennial (Q), and the control (C), which only experienced fire once in 2011. (B) Fire history depicting the months when prescribed bushfires occurred at each plot. Fire severity (thus habitat openness) increases in the following order C < Q < EB < MB < LB.

#### Study species

Here, we show that the three lizard species we studied have varying responses to the fire regimes that each plot was subjected (Fig. S2). *Copeoglossum nigropunctatum* showed fewer captures in intermediate fire severity levels but increased abundance after prescribed fires ended in 2009, while *Micrablepharus atticolus* and *Tropidurus itambere* displayed the opposite pattern (Fig. S2).

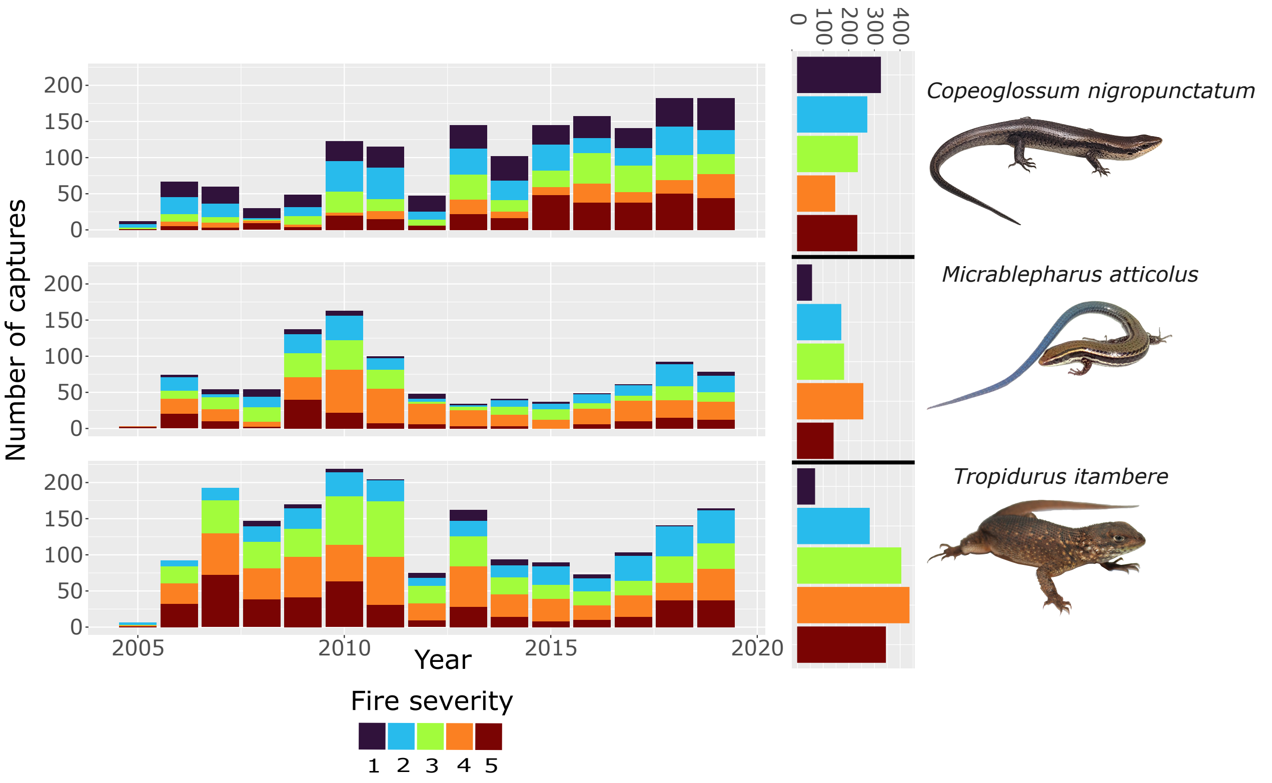


**Fig. S2.** In this study, we investigated the demographic resilience components (resistance, compensation, and recovery time) to different fire timings and frequencies of three lizard species (*Copeoglossum nigropunctatum*, *Micrablepharus atticolus*, and *Tropidurus itambere*) characterized by distinct life-history strategies. Number of captures of three species of lizards (*C. nigropunctatum*, *M. atticolus*, and *T. itambere*) from the Brazilian Cerrado savannas in fire regimes of varying fire severity over the years of study and beside the total captures in each plot. Photos: *C. nigropunctatum* and *M. atticolus* (Nicolás Pelegrin); *T. itambere* (Carlos Morais).

#### Ecophysiological measurements

We estimated thermal performance curves (TPC) and preferred temperature ranges ($t_{pref}$) for each lizard species (Fig. S3). First, to determine the preferred temperatures ($t_{pref}$), we placed the individuals in a ~15–50 °C temperature gradient made of MDF plywood (0.15 m wide × 0.3 m high × 1.0 m long) (Fig. S3A). We placed a 60 W incandescent lamp to generate heat in one corner and ice packs to cool in the other. We left lizards in the gradient for 1 h and recorded their body temperatures every minute with a 1 mm thermocouple attached with tape to their abdomen and connected to a data logger. With the $t_{pref}$ we can estimate the hours of activity based on the microclimate temperature estimates (Fig. S3D). We used the range between the 5th and 95th temperature percentiles as the preferred temperature range for each species (Figs. S3D and S5). Then, we submitted the individuals to different temperatures and stimulated them to run while video recording (Fig. S3B). Individuals ran twice in three temperatures: at ambient temperature (25-28°C), at 5°C below, and at 5°C above said ambient temperature. We recorded the runs with a Casio® EX-FH25 digital camera and calculated their velocity with the Tracker software (https://physlets.org/tracker/). To build the thermal performance curves (TPCs), we also determined the minimum and maximum critical temperatures by submitting individuals to cold and hot temperatures until we observed no righting response (Fig. S3C), *i.e.*, complex muscular movements in response to a stimulus (Taylor *et al.* 2021). Next, we built generalized additive mixed models (GAMMs) with package GAMM4 (Wood & Scheipl 2017) for each species relating maximum speed with body temperature, including individuals as a random factor (Figs. S3E and S4). With the hourly microclimate temperature estimates at 10 cm height, we predicted monthly mean locomotor performance (*perf*) and the hours of activity (*ha*). To predict *perf*, we used the GAMMs for each species. We considered the hours of activity (*ha*) as the hours when microclimate temperatures were within the preferred temperature ranges from each species (Figs. S3 and S5). As the three species are diurnal, we only considered hours of activity when insolation was greater than zero. After the experiments, we returned the individuals to their sites.

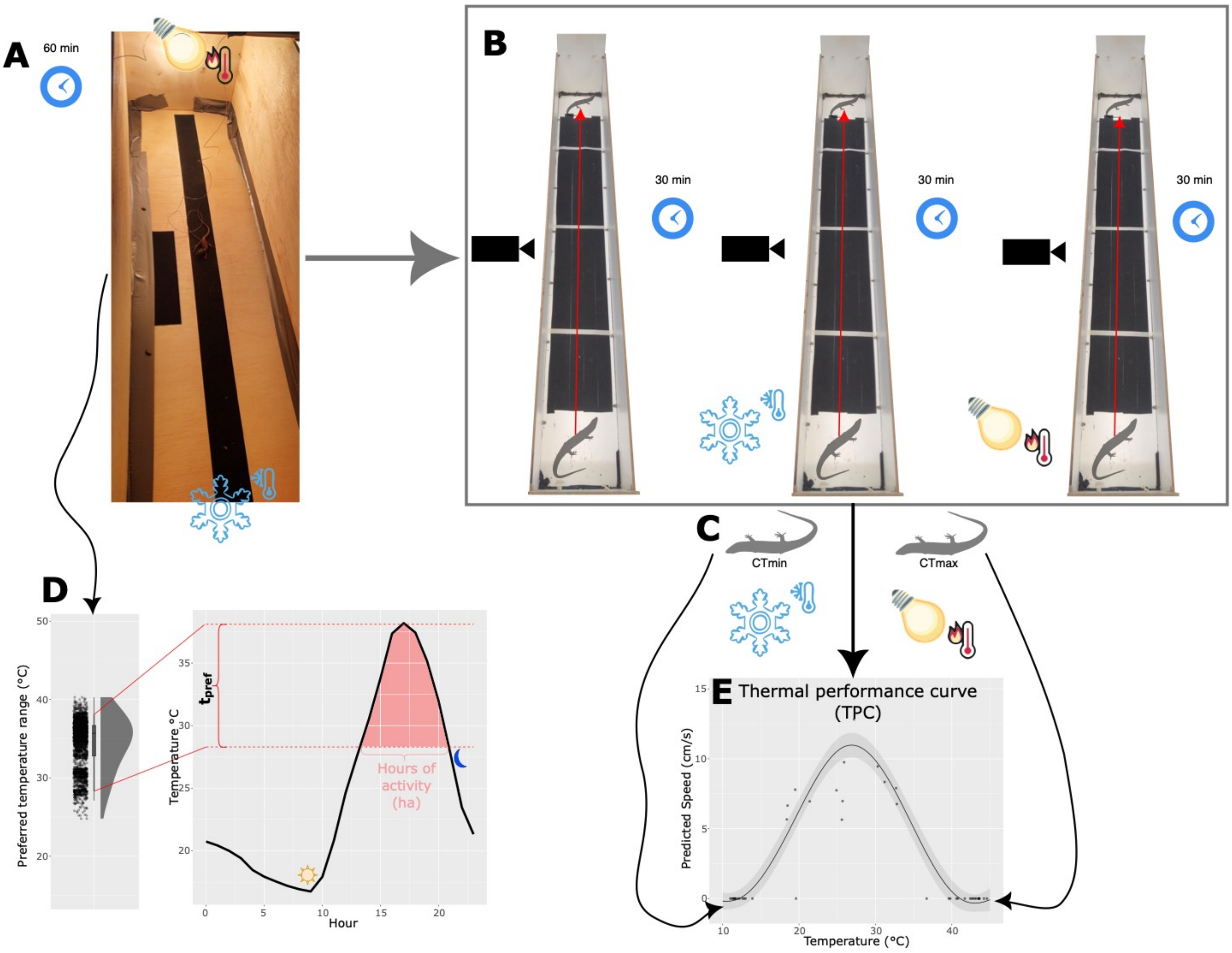


**Fig. S3.** Description of the ecophysiological experiments protocol and examples of the final ecophysiological traits (hours of activity—ha and thermal performance curve—TPC). (A) Measurement of the preferred temperature range in a gradient of ~15-50 °C. Lizards acclimate for 15 min, and we record temperature measures every minute for one hour (60 measures). (B) After, we video-recorded the sprint runs of the lizard in three different temperatures: at ambient temperature (25-28 °C), at 5 °C below, and at 5 °C above said ambient temperature. Lizards rest for 30 min between the runs. (C) Finally, we measured the minimum and maximum critical temperatures ($CT_{min}$ and $CT_{max}$, respectively). (D) We used the range between the 5$^{th}$ and 95$^{th}$ temperature percentiles as the preferred temperature range ($t_{pref}$) for each species. We considered the hours of activity (*ha*) as the hours when microclimate temperatures were within the preferred temperature range for each species. As the three species are diurnal, we only considered hours of activity when insolation was greater than zero. (E) With the maximum speed records from the sprint runs and the critical temperatures ($CT_{min}$ and $CT_{max}$) we built thermal performance curves (TPCs) using generalized additive mixed-effects models.

Despite some differences in the thermal performance curves of the species, all three species have optimal locomotor performances between 25 and 32 °C (Fig. S4).

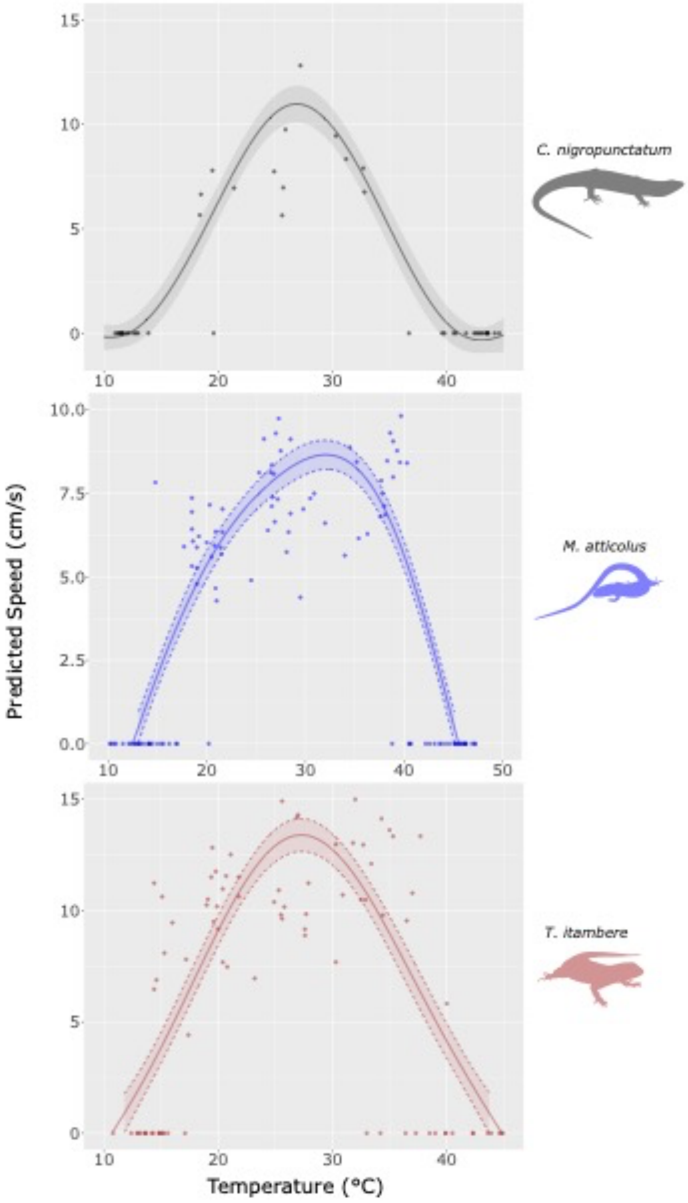


**Fig. S4.** Locomotor thermal performance curves (TPCs) from three species of lizards (*Copeoglossum nigropunctatum*, *Micrablepharus atticolus*, and *Tropidurus itambere*) from the Brazilian Cerrado savannas. Lines and shades represent mean estimates and 95% confidence intervals, respectively.

The species have different thermal preference ranges (5$^{th}$ and 95$^{th}$ temperature percentiles). Preferred temperature of *C. nigropunctatum* ranges between 28.3 and 38.1 °C, *M. atticolus* between 21.0 and 39.4 °C, and *T. itambere* between 24.5 and 40.0 °C (Fig. S5).

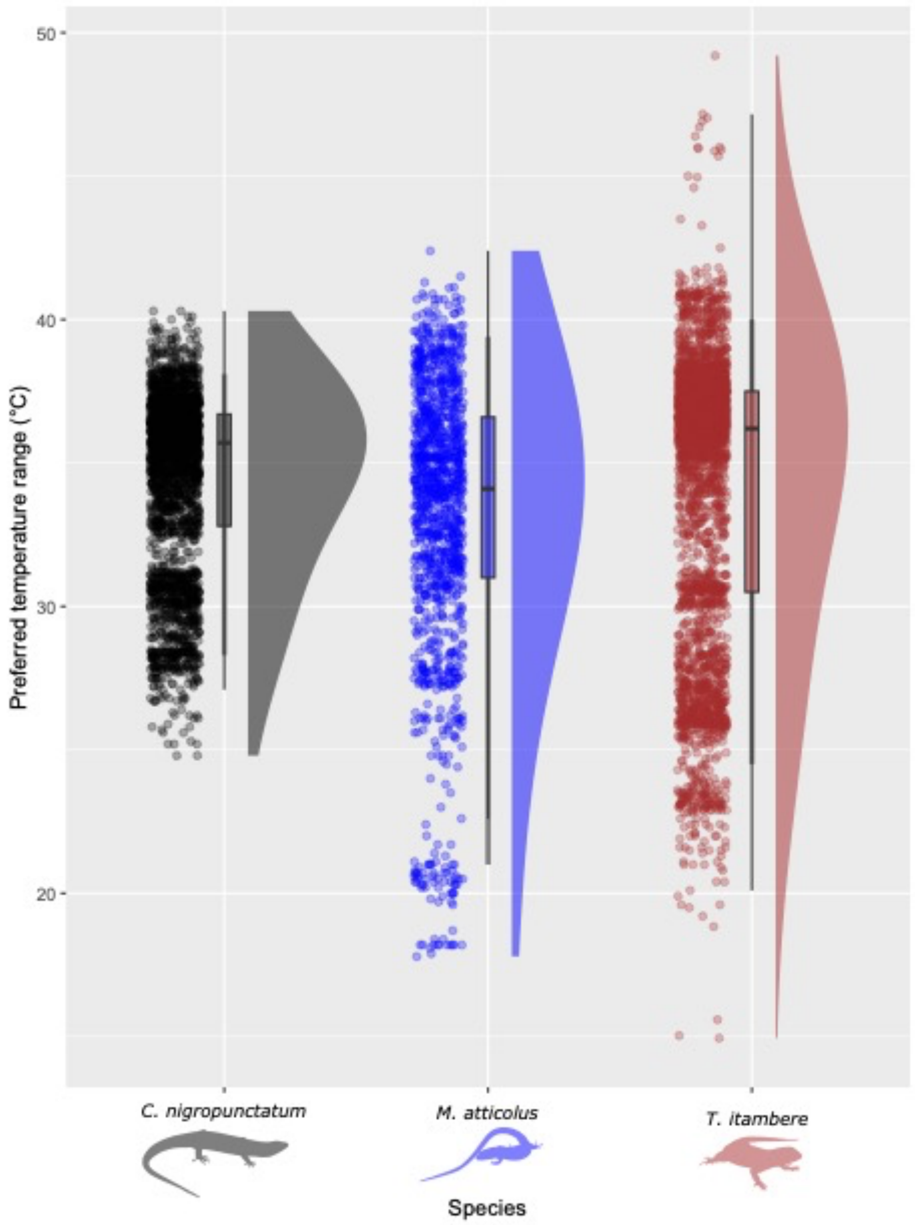


**Fig. S5.** Preferred temperature ranges ($t_{pref}$) from three species of lizards (*Copeoglossum nigropunctatum*, *Micrablepharus atticolus*, and *Tropidurus itambere*) from the Brazilian Cerrado savannas. Boxplots depict median (solid horizontal bars) and interquartile range (boxes). Whiskers extend as far as 1.5x the interquartile range or to minimum and maximum $t_{pref}$. The vertical solid horizontal bars indicate the 90$^{th}$ temperature percentiles.

#### Microclimate estimates

To estimate temperature and air humidity in the different fire regimes, we used a mechanistic model using the ERA-5 dataset from the European Center for Medium-Range Weather Forecasts

(ECMWF) (Hersbach *et al.* 2018) following Kearney *et al.* (2020). To account for differences among fire regimes (plots) and for temporal variation (including vegetation succession) in the microclimate model, we used previous measures of canopy closure from one year of each plot and related them with the leaf area index (LAI) using a generalized linear model with binomial errors (Fig. S6-A). For details on the method of measuring the canopy closure, see (Costa *et al.* 2021). We extracted LAI values of each plot locality from the MODIS product MCD15A2H v006 (Myneni, Knyazikhin & Park 2015) using the package MODISTOOLS. With this model, we predicted the canopy closures for the missing months, ultimately considering the vegetation succession and differences among fire regimes (Fig. S6-B).

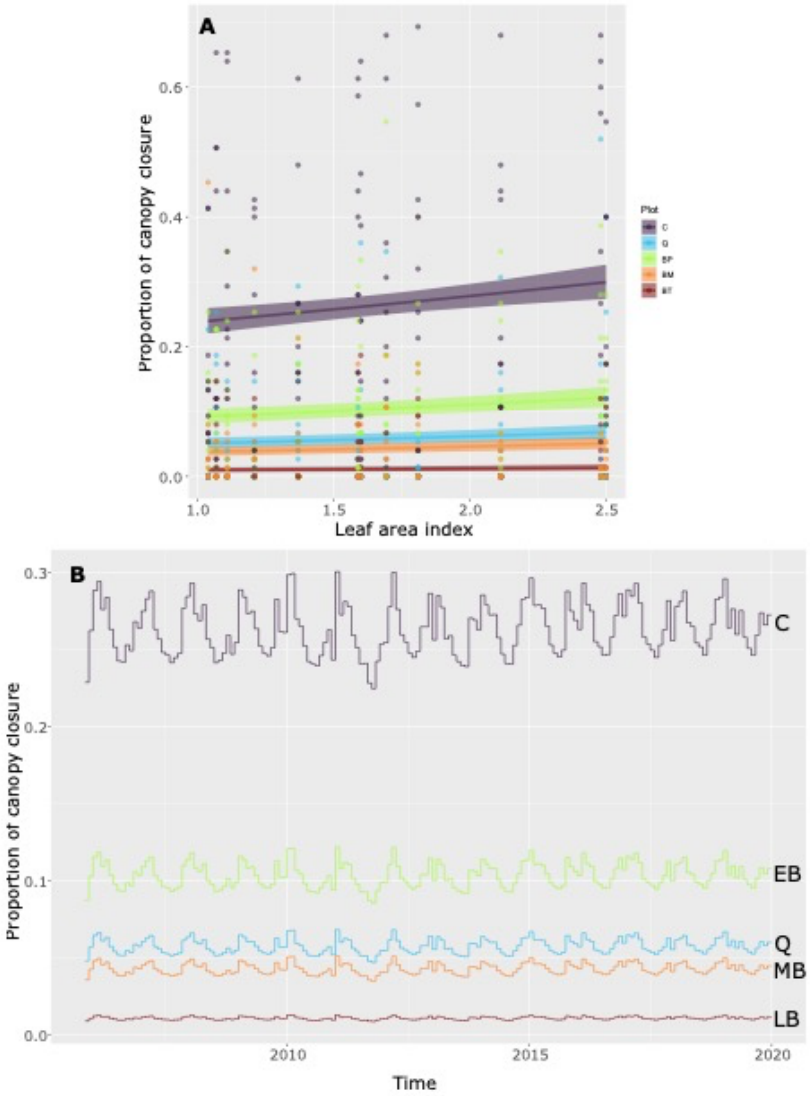


**Fig. S6.** Relationship between the proportion of canopy closure and leaf area index from a generalized linear model (GLM), including differences among plots, each exerted to different fire timings and frequencies (A). Lines and shades represent mean estimates and 95% confidence intervals, respectively. Predictions of canopy closure for the study timeframe (November 2005 to December 2019) using the GLM and leaf area index values. C = control, Q = quadrennial, EB = early biennial, MB = mid biennial, LB = late biennial.

#### Models' parameterizations

First, we compared and selected models using the Deviance Information Criteria (DIC) (Spiegelhalter *et al.* 2002) with linear and quadratic relationships between the individuals' snout-vent length (SVL and SVL + SVL$^2$) with the vital rates (survival, growth, probability of reproduction, and number of newborns) (Table S1).

**Table S1.** Models' comparison using Deviance Information Criteria (DIC) to assess the performance of linear and quadratic relationships between the individuals' snout-vent length (SVL) and the vital rates (survival, growth, probability of reproduction–$p_{rep}$, and number of newborns–$n_b$) of three lizard species from the Brazilian Cerrado savannas. We highlighted with bold letters the lowest DIC.

| Species | Vital rate | SVL | SVL + SVL$^2$ |
|---|---|---|---|
| *C. nigropunctatum* | CJS / Von Bertalanffy (Survival/Growth) | 71718.11 | **68525.63** |
| | $p_{rep}$ | 171.08 | **170.73** |
| | $n_b$ | **425.61** | 428.07 |
| *M. atticolus* | CJS / Von Bertalanffy (Survival/Growth) | **34511.86** | 56410.47 |
| | $p_{rep}$ | **230.97** | 231.69 |
| | $n_b$ | - | - |
| *T. itambere* | CJS / Von Bertalanffy (Survival/Growth) | **131779.40** | 196867.00 |
| | $p_{rep}$ | **100.23** | 101.06 |
| | $n_b$ | 203.58 | **202.98** |

The Pradel Jolly-Sebber (PJS) model is a time-symmetric open model that estimates survival ($\sigma$) and recruitment per-capita rates (*f*) (Pradel 1996). Our PJS model included the environmental stochasticity from the microclimate, weather, ecophysiological traits (locomotor performance–*perf* and hours of activity–*ha*), fire regime (*plot*), fire occurrence (month with fire = 1; month with no fire = 0) and time since the last fire (*TSLF*, in months) into the survival and recruitment estimates. Microclimatic and ecophysiological variables varied among plots, as we explained in the previous section. We present the correlations and relationships among the variables in Fig. S7. The following equation notations group all the environmental variables (weather, microclimate, and ecophysiological) to only *env*.

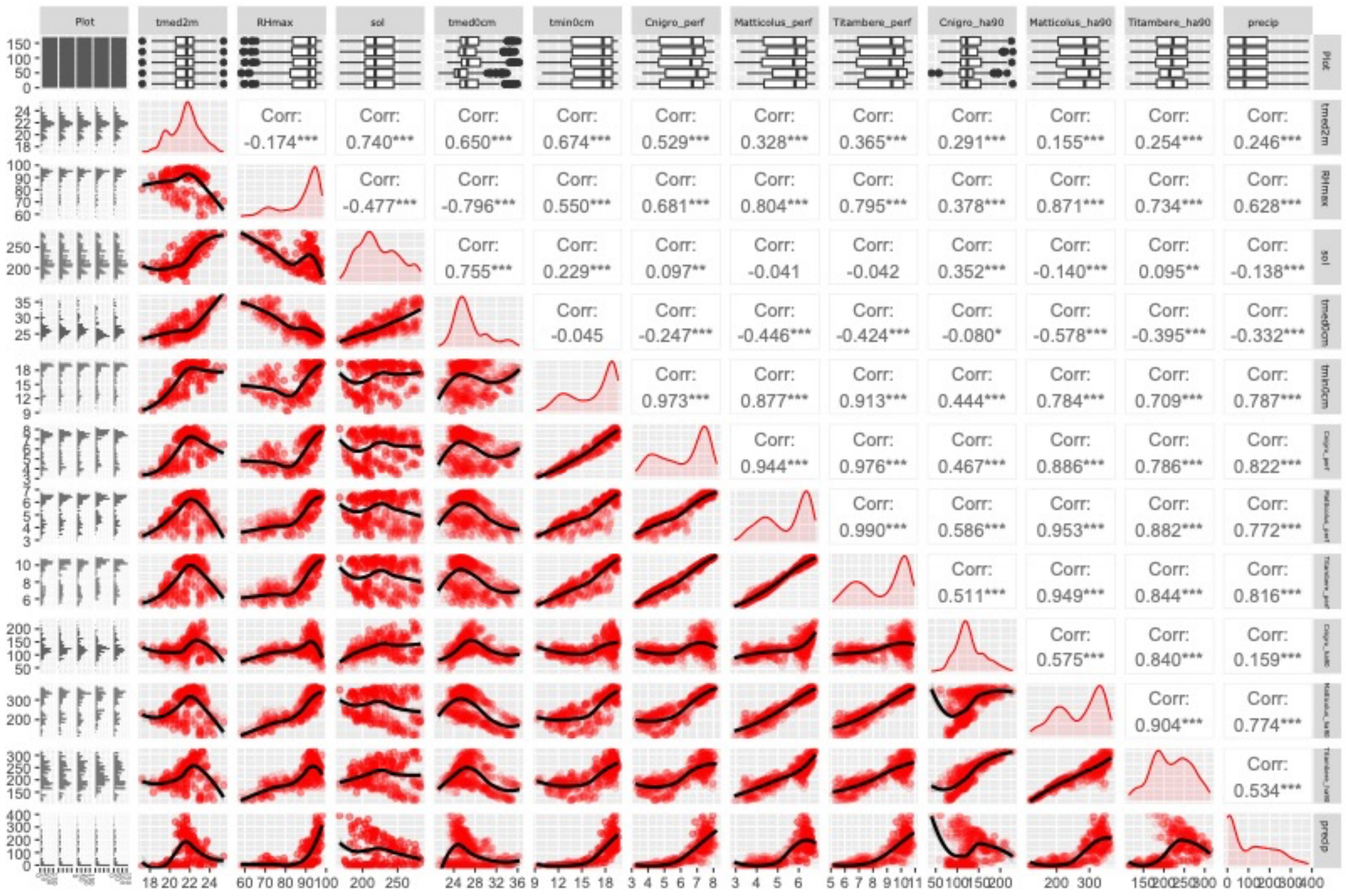


**Fig. S7.** Correlations between monthly weather/microclimatic/ecophysiological variables used in the mark-recapture models to estimate the effects of environmental covariates on vital rates of three species of lizards (*Copeoglossum nigropunctatum*, *Micrablepharus atticolus*, and *Tropidurus itambere*) from the Brazilian Cerrado savannas in fire regimes of varying fire severity. *tmed2m* = mean air temperature at 200 cm height; *tmin0cm* = minimum air temperature at 0 cm height; *tmed0cm* = mean air temperature at 0 cm height; *RHmax* = maximum relative air humidity; *sol* = solar radiation; *precip* = accumulated precipitation; *perf* = mean locomotor performance (one for each species); *ha* = hours of activity (one for each species).

To account for unexplained stochastic temporal variation in the populations' survival estimates, we also included a monthly random variation $\epsilon(plot, t)$. In summary, the populations' survival ($\overline{\sigma_{PJS}(plot, t)}$) was modeled as

$$logit\left(\overline{\sigma_{PJS}(plot, t)}\right) = \alpha_{\sigma}(plot) + \beta_{\sigma env} \cdot env(plot, t) + \epsilon_{\sigma}(plot, t) \quad \text{(Eq. S1)}$$

where each plot had its own intercept ($\alpha_{\sigma}(plot)$) and all environmental predictors were included (summarized in the term $\beta_{\sigma env} \cdot env(plot, t)$). To estimate the variable importance for each environmental predictor we used the indicator variable selection, where each indicator had a Bernoulli prior probability of 0.5 (Kuo & Mallick 1998; O'Hara & Sillanpää 2009). Thus, we model-averaged the covariate coefficients estimates ($\beta_{\sigma env}$) by averaging across posterior samples conditional on the covariate being present in the model (indicator variable selection = 1) (Royle & Dorazio 2008).

The Cormack-Jolly-Seber (CJS) model (eq. S2) related the individuals' survival with the individuals' *SVL* as:

$$logit\left(\sigma_{CJS}(SVL)\right) = \overline{\sigma_{PJS}(plot, t)} + \beta_{SVL} \cdot SVL + \beta_{SVL^2} \cdot SVL^2 \quad \text{(Eq. S2)}$$

where the populations' survival estimate ($\overline{\sigma_{PJS}(plot, t)}$) was the intercept of the model. The quadratic term ($\beta_{SVL_{it}^2} \cdot SVL_{it}^2$) was (or was not) included based on the previous models' comparisons using DIC (Table S1). We used the same approach for capture probability parameters ($p_{CJS}$ for individuals' capture probability and $\overline{p_{PJS}}$ for populations' capture probability).

For the transition of the body size through time (growth, $\gamma$), we combined a Von Bertalanffy body growth function with the individuals' records of SVL and capture history (eq. S3). The body size of individual $i$ at time $t$ ($SVL_{it}$) is determined by:

$$SVL_t = SVL0 + (SVLI - SVL0) \cdot (1 - K(plot)^{(AFC + \Delta t)}) \text{ (Eq. S3)}$$

where $AFC$ is the age of the individual at first capture, and $\Delta t$ is the number of months since first capture (Schofield, Barker & Taylor 2013; Reinke *et al.* 2020). The sum of these parameters is the age of the individual $i$ at time $t$. The initial size at age 0 for individual $i$ is denoted as $SVL0$, the asymptotic size as $SVLI$, and $K$ is the growth constant (measured in amounts of time to reach asymptotic size). All variables were allowed to vary through individuals, but $K$ also varied with different means among the plots.

For reproduction, we considered three different processes: the probability of reproduction ($p_{rep}$), the production of newborns (clutch size; $n_b$), and the probability of recruitment ($p_{rec}$). To estimate the probability of reproduction, we used the records of reproductive and non-reproductive females we captured in the field ($y_{prep}$) and related with the individual's $SVL$ using a Bernoulli distribution as follow:

$$y_{prep} \sim Bernoulli(p_{rep}),\ logit(p_{rep}) = m_0 + \beta_{prepSVL} \cdot SVL + \beta_{prepSVL^2} \cdot SVL^2 \text{ (Eq. S4)}$$

where $m_0$ is the intercept and $\beta_{prepSVL}$ is the angular coefficient of the relationship. The quadratic term ($\beta_{prepSVL^2} \cdot SVL^2$) was (or was not) included based on the previous models' comparisons using DIC (Table S1).

To estimate the production of newborns $n_b$, we used a Poisson distribution relating the number of eggs or embryos produced per female ($y_{n_b}$) to the individual's $SVL$ as follow:

$$y_{n_b} \sim Poisson(n_b),\ log(n_b) = n_0 + \beta_{nbSVL} \cdot SVL + \beta_{nbSVL^2} \cdot SVL^2 \quad \text{(Eq. S5)}$$

where $n_0$ is the intercept and $\beta_{nbSVL}$ is the angular coefficient of the relationship. The quadratic term ($\beta_{nbSVL^2} \cdot SVL^2$) was (or was not) included based on the previous models' comparisons using DIC (Table S1). Due to logistical limitations, data on the number of eggs or embryos came from *ex-situ* locations, mainly specimens deposited in the Herpetological Collection of the University of Brasília (CHUNB).

We estimated the *per capita* recruitment (*f*) using the PJS model with the same environmental predictors used for survival and capture populations' estimates ($\overline{\sigma_{PJS}}$ and $\overline{p_{PJS}}$, respectively), as the following equation:

$$log(f(plot,t)) = \alpha_f(plot) + \beta_{fenv} \cdot env(plot,t) + \epsilon_f(plot,t) \quad \text{(Eq. S6)}$$

where each plot had its own intercept ($\alpha_f(plot)$) and all environmental predictors included (summarized in the term $\beta_{fenv} \cdot env(plot,t)$). We performed the variable selection of the environmental predictors and model-averaging with the same approach of the other PJS parameters ($\overline{\sigma_{PJS}}$ and $\overline{p_{PJS}}$). We also included a monthly random variation $\epsilon_f(plot,t)$ to account for unexplained stochastic temporal variation. Then, following Pradel (1996), we derived the probability of recruitment ($p_{rec}$) as:

$$p_{rec}(plot,t) = 1 - \frac{f(plot,t)}{\left(\overline{\sigma_{PJS}(plot,t)} + f(plot,t)\right)} \quad \text{(Eq. S7).}$$

In our IPMs, The $P$ sub-kernel incorporates both survival and growth components, such that $P(x,y) = \sigma(x) \cdot \gamma(x,y)$, where $\sigma(x)$ is the survival probability of a size-$x$ individual and $\gamma(x, y)$ is the probability of a size-$x$ individual growing to size $y$. The survival probability was determined by the $logit\left(\sigma_{CJS}(x)\right)$ and the growth probability function by $\gamma(x,y) = f_N(\overline{SVL(x)}, \tau^2(x))$, where $f_N$ is a normal probability density, $\overline{SVL}$ is the expected mean of SVL and $\tau^2$ is the expected standard deviation of $SVL$, given the size-$x$ individual and the estimate samples of the Von Bertalanffy function from the hierarchical Bayesian model.

The $F$ sub-kernel was determined by:

$$F(x,y) = \left(p_{rec}(plot,t) + p_{rep}(SVL)\right) \cdot n_b(SVL) \cdot dSVL_{nb} \qquad \text{(Eq. S8)}$$

where $dSVL_{nb}$ is the normal size distribution of newborns, with mean and standard deviations measured with the newborn individuals we captured in the field. Thus, we considered the fecundity sub-kernel as the sum of the probabilities of recruitment and reproduction, multiplied by the production of newborns and their size distribution. The probability of recruitment is the only parameter that incorporates the environmental dependency into the reproduction portion of the IPM.

We note that CJS and PJS models cannot differentiate between permanent emigration or mortality and immigration or fecundity when estimating survival and recruitment, respectively (Cormack 1964; Pradel 1996). However, our data show that peaks of the recruitment estimates coincide with the captures of hatchlings in the three species, reinforcing our modeling approach (Fig. S8). Permanent emigration of individuals has the same implication of individuals' mortality for the population size (both are exits of individuals out of the population), thus the "apparent survival" estimates of CJS and PJS models do not compromise the interpretation of our results. Moreover, the nature of our hypotheses and available data do not justify a more complex model considering migration between plots, since the lizard species of our study have low dispersal potential (Vitt &

Blackburn 1991; Rodrigues 1996; Van Sluys 1997; Vitt, Zani & Lima 1997; Galán 1999; Vieira et al. 2000), and we did not observe individuals recaptured in different plots in all the study period (pers. obs.).

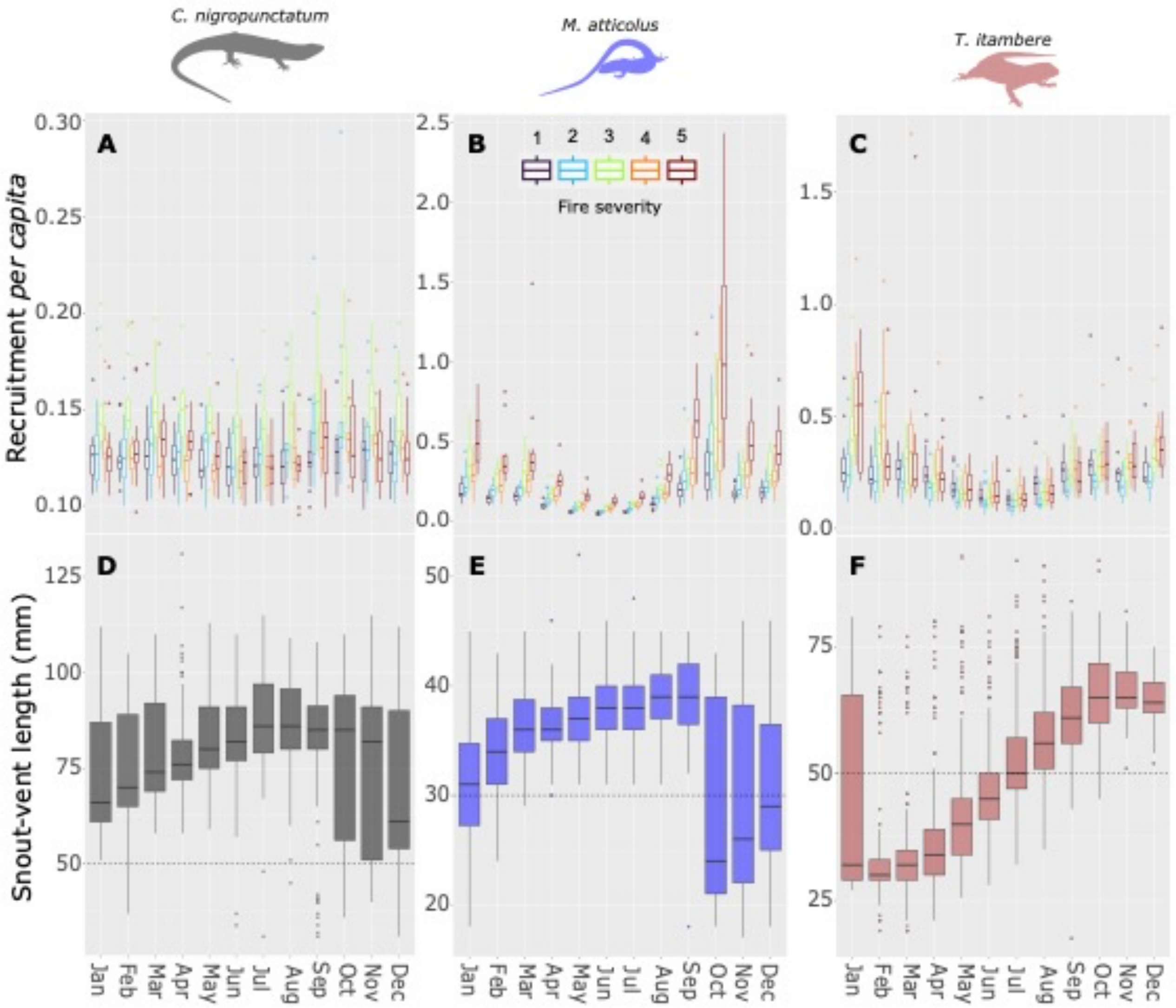


**Fig. S8.** Variation of recruitment *per capita* (A, B, and C) and snout-vent length (D, E, and F) of populations of three lizard species (*Copeoglossum nigropunctatum*–A and D, *Micrablepharus atticolus* – B and E, and *Tropidurus itambere*–C and F) from the Brazilian Cerrado savannas in fire regimes of varying severity. Note that peaks of recruitment estimates coincide with small individuals (hatchlings).

We implemented the models for each species with JAGS in R, using the packages JAGSUI, RUNJAGS, and RJAGS (Kellner 2019; Plummer 2019). We initially used four Markov chains of 50,000 iterations in the JAGS adaptive phase, discarded 50,000 iterations as burn-in, and sampled 10,000 estimates by a thinning rate of 10. We assessed convergence within and between MCMC sequences with trace and density plots and the potential scale reduction factor ($\hat{R}$), considering that convergence was satisfactorily approached when $\hat{R}$ was smaller than 1.1 for all parameters (Gelman *et al.* 2014). However, we experienced convergence issues with some parameters in the CJS and PJS models of *C. nigropunctatum* (specifically, for $\overline{\sigma_{PJS}}$) and *T. itambere* (for $f$). We ran these models with different parameterizations and MCMC settings to improve convergence (see vital rates estimates section below).

**Sex-analyses**

We only began to register the sex of individuals in 2010. We have not found sex differences in survival and growth (Fig. S9) to justify using female-centred IPMs.

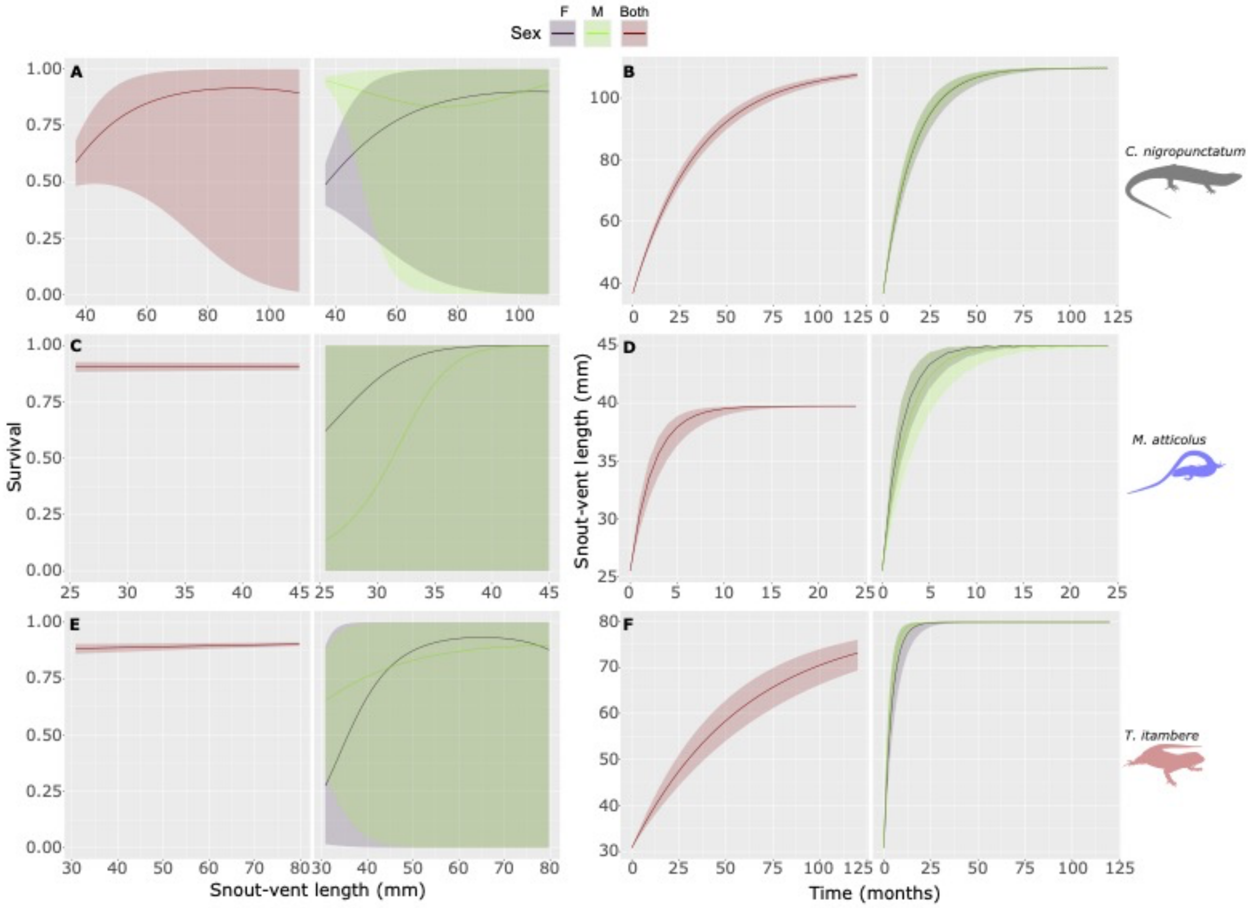


**Fig. S9.** Estimates of survival (predicted by individuals' body size – A, C & E) and body growth rates (B, D & F) of three lizard species (A-B) *Copeoglossum nigropunctatum*, (C-D) *Micrablepharus atticolus*, and (E-F) *Tropidurus itambere* under fire regimes of varying severity. Lines and shades represent mean estimates and 95% credible intervals derived from a joint Bayesian Cormack-Jolly Seber (CJS) mark-recapture (survival) and Von Bertalanffy growth models. The sex-based models use data between 2010 and 2019, while the sex-averaged model (both sexes) is between 2005 and 2019.

#### Brute force sensitivity analysis

To implement the brute force sensitivity analysis, we added values from 0.01 to 0.1 to each of the parameters specifying our vital rates and recalculated the differences in the demographic resilience components. Our parameter sensitivities ($S_p$) were estimated as in equation S9, where the numerator describes the difference in the transient metric of choice (*e.g.*, reactivity for amplification) resulting from perturbing a vital rate parameter *p* at a time, in the denominator, following Cant *et al.* (2023).

$$S_p = \frac{Transient\ metric_{\ original} - Transient\ metric_{perturbed}}{p_{original} - p_{perturbed}} \text{ (Eq. S9)}$$

**Mark-recapture descriptive statistics**

Over the 14 years, we made 1,557 captures from 1,207 individuals of *Copeoglossum nigropunctatum*, 1,025 from 803 individuals of *Micrablepharus atticolus*, and 1,932 from 1528 individuals of *Tropidurus itambere*. The maximum number of recaptures between months for one individual was four for *C. nigropunctatum*, six for *M. atticolus*, and five for *T. itambere*. On average, individuals were recaptured through 7.82 ± 9.01 SD months in *C. nigropunctatum*, 6.00 ± 8.99 SD in *M. atticolus*, and 11.30 ± 22.83 SD in *T. itambere*. The maximum period between the first and last recapture of one individual was 55 months in *C. nigropunctatum*, 40 months in *M. atticolus*, and 38 months in *T. itambere*.

**Vital rate estimates**

In this section, we present the results and diagnostics from the Bayesian hierarchical models of the vital rates we estimated. When some parameters have not converged, we ran each model separately, increased the number of iterations, or reparametrized it.

*Survival/Growth models*

The CJS and Von Bertalanffy integrated models converged for *M. atticolus* and *T. itambere* with the initial MCMC settings, but not for *C. nigropunctatum*. Therefore, we ran the CJS model separately for *C. nigropunctatum* with four Markov chains of 50,000 iterations in the JAGS adaptive phase, discarded 100,000 iterations as burn-in, and sampled 500,000 estimates with no thinning. Table S2 presents the results of the CJS/Von Bertalanffy models for the three species.

**Table S2.** Cormack-Jolly-Seber (CJS) and Von Bertalanffy model estimates (*Mean* ± *SD*) and credible interval (2.5% and 97.5%) of the relationship between individuals' snout-vent length (SVL) and survival (σ), capture (*p*), and body growth parameters of three lizard species from the Brazilian Cerrado savannas. Survival and capture probabilities and body growth constants are in the logit scale. Values of $\hat{R}$ closer to 1.00 indicate model convergence. α = intercept; μ = mean; K = body growth constant; SVLI = asymptotic size.

| **Parameter** | **Vital rate** | **Mean** | **SD** | **2.5%** | **97.5%** | $\hat{R}$ | **Effective sample size** |
|---|---|---|---|---|---|---|---|
| | | | ***Copeoglossum nigropunctatum*** | | | | |
| $\alpha_{\sigma}$ | Individuals' survival | -3.310 | 1.478 | -6.233 | -0.483 | 1.081 | 148 |
| $\beta_{\sigma SVL}$ | Individuals' survival | 0.125 | 0.042 | 0.047 | 0.208 | 1.087 | 106 |
| $\beta_{\sigma SVL^2}$ | Individuals' survival | -0.001 | 0.000 | -0.001 | 0.000 | 1.089 | 119 |
| $\alpha_{p}$ | Individuals' capture | -2.751 | 1.899 | -6.364 | 0.986 | 1.066 | 199 |
| $\beta_{pSVL}$ | Individuals' capture | 0.013 | 0.054 | -0.094 | 0.113 | 1.066 | 172 |
| $\beta_{pSVL^2}$ | Individuals' capture | 0.000 | 0.000 | -0.001 | 0.001 | 1.064 | 182 |
| $\mu_{K}(C)$ | Individuals' body growth | 3.640 | 0.042 | 3.557 | 3.722 | 1.000 | 2690 |

| Parameter | Vital rate | Mean | SD | 2.5% | 97.5% | $\hat{R}$ | Effective sample size |
|---|---|---|---|---|---|---|---|
| $\mu_K$(Q) | Individuals' body growth | 3.799 | 0.039 | 3.720 | 3.873 | 1.001 | 3961 |
| $\mu_K$(EB) | Individuals' body growth | 3.465 | 0.051 | 3.364 | 3.562 | 1.001 | 3259 |
| $\mu_K$(MB) | Individuals' body growth | 3.293 | 0.072 | 3.154 | 3.434 | 1.001 | 3267 |
| $\mu_K$(LB) | Individuals' body growth | 3.522 | 0.049 | 3.426 | 3.618 | 1.000 | 4111 |
| $\mu_{SVLI}$ | Individuals' body growth | 109.953 | 0.223 | 109.515 | 110.381 | 1.002 | 4623 |
| | | | ***Micrablepharus atticolus*** | | | | |
| $\beta_{\sigma SVL}$ | Individual survival | 0.000 | 0.004 | -0.007 | 0.008 | 1.020 | 693 |
| $\beta_{pSVL}$ | Individual capture | -0.002 | 0.004 | -0.010 | 0.006 | 1.000 | 898 |
| $\mu_K$(C) | Individual Body growth | 0.564 | 0.323 | 0.042 | 1.239 | 1.010 | 2059 |
| $\mu_K$(Q) | Individual Body growth | 1.152 | 0.230 | 0.679 | 1.579 | 1.020 | 1217 |
| $\mu_K$(EB) | Individual Body growth | 0.447 | 0.229 | 0.048 | 0.909 | 1.020 | 801 |

| Parameter | Vital rate | Mean | SD | 2.5% | 97.5% | $\hat{R}$ | Effective sample size |
|---|---|---|---|---|---|---|---|
| $\mu_K$(MB) | Individual Body growth | 0.084 | 0.076 | 0.002 | 0.284 | 1.000 | 2147 |
| $\mu_K$(LB) | Individual Body growth | 1.110 | 0.211 | 0.673 | 1.504 | 1.010 | 1890 |
| $\mu_{SVLI}$ | Individual Body growth | 39.731 | 0.212 | 39.354 | 40.176 | 1.070 | 323 |
| | | | ***Tropidurus itambere*** | | | | |
| $\beta_{\sigma SVL}$ | Individual survival | 0.004 | 0.002 | 0.000 | 0.009 | 1.050 | 857 |
| $\beta_{pSVL}$ | Individual capture | -0.004 | 0.003 | -0.009 | 0.001 | 1.030 | 1706 |
| $\mu_K$(C) | Individual body growth | 4.340 | 0.222 | 3.909 | 4.780 | 1.000 | 3783 |
| $\mu_K$(Q) | Individual body growth | 4.467 | 0.125 | 4.220 | 4.716 | 1.030 | 2794 |
| $\mu_K$(EB) | Individual body growth | 4.271 | 0.096 | 4.083 | 4.463 | 1.000 | 1577 |
| $\mu_K$(MB) | Individual body growth | 3.046 | 0.096 | 2.861 | 3.237 | 1.030 | 1038 |
| $\mu_K$(LB) | Individual body growth | 4.329 | 0.106 | 4.125 | 4.537 | 1.030 | 1406 |

| Parameter | Vital rate | Mean | SD | 2.5% | 97.5% | $\hat{R}$ | Effective sample size |
|---|---|---|---|---|---|---|---|
| $\mu_{SVLI}$ | Individual body growth | 79.814 | 0.222 | 79.383 | 80.247 | 1.000 | 2026 |

*Probability of reproduction and production of newborns*

The parameters from the generalized linear models of the $p_{rep}$ and $n_b$ converged with the initial MCMC settings, except for the $n_b$ in *T. itambere*. For this vital rate, we ran four Markov chains of 100,000 iterations in the JAGS adaptive phase, discarded 1,800,000 iterations as burn-in, and sampled 200,000 estimates by a thinning rate of 100. Table S3 presents the results of the CJS/Von Bertalanffy models for the three species.

**Table S3.** Generalized linear models estimates (*Mean* ± *SD*) and credible interval (2.5% and 97.5%) of the relationship between individuals' snout-vent length (SVL) and number of newborns ($n_b$) and probability of reproduction ($p_{rep}$) of three lizard species from the Brazilian Cerrado savannas. The probability of reproduction is in the logit scale, and the number of newborns is in the log scale. Values of $\hat{R}$ closer to 1.00 indicate model convergence. α = intercept coefficient; β = slope coefficient.

| Parameter | Vital rate | Mean | SD | 2.5% | 97.5% | $\hat{R}$ | Effective sample size |
|---|---|---|---|---|---|---|---|
| ***Copeoglossum nigropunctatum*** | | | | | | | |
| $\alpha_{nb}$ | Number of newborns | -0.792 | 0.493 | -1.739 | 0.204 | 1.002 | 969 |

| Parameter | Vital rate | Mean | SD | 2.5% | 97.5% | $\hat{R}$ | Effective sample size |
|---|---|---|---|---|---|---|---|
| $\beta_{nbSVL}$ | Number of newborns | 0.025 | 0.005 | 0.015 | 0.035 | 1.002 | 973 |
| $\alpha_{prep}$ | Probability of reproduction | -6.392 | 2.858 | -10.000 | -0.900 | 1.010 | 208 |
| $\beta_{prepSVL}$ | Probability of reproduction | 0.014 | 0.066 | -0.116 | 0.117 | 1.010 | 201 |
| $\beta_{prepSVL}^2$ | Probability of reproduction | 0.000 | 0.000 | 0.000 | 0.001 | 1.009 | 246 |
| | | ***Micrablepharus atticolus*** | | | | | |
| $\alpha_{prep}$ | Probability of reproduction | -8.139 | 1.385 | -9.926 | -4.875 | 1.000 | 1656 |
| $\beta_{prepSVL}$ | Probability of reproduction | 0.151 | 0.034 | 0.070 | 0.196 | 1.000 | 1652 |
| | | ***Tropidurus itambere*** | | | | | |
| $\alpha_{nb}$ | Number of newborns | -11.299 | 6.804 | -24.712 | 1.910 | 1.000 | 40416 |
| $\beta_{nbSVL}$ | Number of newborns | 0.310 | 0.199 | -0.080 | 0.696 | 1.000 | 40485 |
| $\beta_{nbSVL}^2$ | Number of newborns | -0.002 | 0.001 | -0.005 | 0.001 | 1.000 | 40552 |
| $\alpha_{prep}$ | Probability of reproduction | -8.194 | 1.277 | -9.920 | -5.208 | 1.000 | 5111 |

| **Parameter** | **Vital rate** | **Mean** | **SD** | **2.5%** | **97.5%** | $\hat{R}$ | **Effective sample size** |
|---|---|---|---|---|---|---|---|
| $\beta_{prepSVL}$ | Probability of reproduction | 0.087 | 0.020 | 0.040 | 0.116 | 1.000 | 5159 |

*Pradel Jolly-Seber models*

The PJS model only converged with the initial MCMC settings for *M. atticolus*. For *C. nigropunctatum*, we reran the model with four Markov chains of 50,000 iterations in the JAGS adaptive phase, discarded 200,000 iterations as burn-in, and sampled 100,000 estimates with no thinning to achieve convergence. For the recruitment rate of *T. itambere*, we had bad mixing for the parameters $\beta_{ftmin0cm}$ and $\beta_{fperf}$, probably related to collinearity with other two important climatic variables: solar radiation (*sol*) and mean air temperature at 200 cm (*tmed2m*). The parameters $\beta_{fftmin0cm}$ and $\beta_{fperf}$ also probably confound the effects of the fire regimes ($\alpha_{f(plot)}$). Therefore, we fixed the importance of the climatic and microclimatic variables (Table S4). We reran the reparametrized model with four Markov chains of 50,000 iterations in the JAGS adaptive phase, discarded 400,000 iterations as burn-in, and sampled 100,000 estimates with no thinning to achieve convergence.

**Table S4.** Model-averaged estimates (*β - Mean* ± *SD*) and credible interval (2.5% and 97.5%) of the demographic parameters – survival (σ), capture (*p*), and recruitment (*f*) – of three lizard species from the Brazilian Cerrado savannas. Survival and capture probabilities are in the logit scale, while recruitment is in the log scale. Values of $\hat{R}$ closer to 1.00 indicate model convergence. Importance values greater than 0.5 (prior) indicate a higher degree of importance in predicting the demographic parameter. α = intercept; ϵ = monthly random variation; *tmed2m* = mean air temperature at 200 cm height; *tmin0cm* = minimum air temperature at 0 cm height; *tmed0cm* = mean air temperature at 0 cm height; *RHmax* = maximum relative air humidity; *sol* = solar radiation; *precip* = accumulated precipitation; *perf* = mean locomotor performance; *ha* = hours of activity; *fire* = fire occurrence; *TSLF* = time since last fire.

| Parameter | Vital rate | Mean | SD | 2.50% | 97.50% | $\hat{R}$ | Effective sample size | Mean importance | SD importance |
|---|---|---|---|---|---|---|---|---|---|
| | | | | ***Copeoglossum nigropunctatum*** | | | | | |
| $\alpha_{\sigma(C)}$ | Population survival | 2.421 | 0.266 | 1.983 | 3.012 | 1.020 | 345 | - | - |
| $\alpha_{\sigma(Q)}$ | Population survival | 2.412 | 0.250 | 1.999 | 2.970 | 1.020 | 476 | - | - |
| $\alpha_{\sigma(EB)}$ | Population survival | 2.294 | 0.266 | 1.847 | 2.868 | 1.010 | 492 | - | - |
| $\alpha_{\sigma(MB)}$ | Population survival | 2.512 | 0.344 | 1.900 | 3.236 | 1.010 | 525 | - | - |
| $\alpha_{\sigma(LB)}$ | Population survival | 2.580 | 0.261 | 2.142 | 3.154 | 1.020 | 740 | - | - |
| $\epsilon_{\sigma(plot,t)}$ | Population survival | 0.852 | 0.300 | 0.209 | 1.436 | 1.020 | 194 | - | - |
| $\beta_{\sigma tmed2m}$ | Population survival | 0.015 | 0.079 | -0.062 | 0.177 | 1.040 | 1950 | 0.503 | 0.500 |

| Parameter | Vital rate | Mean | SD | 2.50% | 97.50% | $\widehat{R}$ | Effective sample size | Mean importance | SD importance |
|---|---|---|---|---|---|---|---|---|---|
| $\beta_{\sigma RHmax}$ | Population survival | 0.011 | 0.049 | -0.058 | 0.138 | 1.000 | 8613 | 0.508 | 0.500 |
| $\beta_{\sigma sol}$ | Population survival | -0.038 | 0.131 | -0.434 | 0.046 | 1.020 | 801 | 0.548 | 0.498 |
| $\beta_{\sigma tmed0cm}$ | Population survival | -0.005 | 0.043 | -0.106 | 0.071 | 1.000 | 9968 | 0.492 | 0.500 |
| $\beta_{tmin0cm}$ | Population survival | 0.017 | 0.070 | -0.054 | 0.186 | 1.000 | 3027 | 0.515 | 0.500 |
| $\beta_{\sigma precip}$ | Population survival | -0.006 | 0.067 | -0.127 | 0.081 | 1.000 | 3083 | 0.496 | 0.500 |
| $\beta_{\sigma perf}$ | Population survival | 0.013 | 0.061 | -0.059 | 0.166 | 1.010 | 4857 | 0.510 | 0.500 |
| $\beta_{\sigma ha}$ | Population survival | 0.004 | 0.040 | -0.068 | 0.098 | 1.000 | 21231 | 0.483 | 0.500 |
| $\beta_{\sigma fire}$ | Population survival | -0.003 | 0.070 | -0.116 | 0.094 | 1.000 | 28086 | 0.501 | 0.500 |
| $\beta_{\sigma TSLF}$ | Population survival | 0.004 | 0.026 | -0.043 | 0.072 | 1.000 | 8592 | 0.440 | 0.496 |
| $\alpha_{f(C)}$ | Population recruitment | -2.290 | 0.189 | -2.703 | -1.963 | 1.010 | 415 | - | - |
| $\alpha_{f(Q)}$ | Population recruitment | -2.255 | 0.179 | -2.675 | -1.960 | 1.010 | 543 | - | - |
| $\alpha_{f(EB)}$ | Population recruitment | -2.127 | 0.182 | -2.520 | -1.812 | 1.010 | 563 | - | - |
| $\alpha_{f(MB)}$ | Population recruitment | -2.311 | 0.258 | -2.855 | -1.851 | 1.000 | 588 | - | - |

| Parameter | Vital rate | Mean | SD | 2.50% | 97.50% | $\widehat{R}$ | Effective sample size | Mean importance | SD importance |
|---|---|---|---|---|---|---|---|---|---|
| $\alpha_{f(LB)}$ | Population recruitment | -2.293 | 0.166 | -2.651 | -2.003 | 1.010 | 1057 | - | - |
| $\epsilon_{f(plot,t)}$ | Population recruitment | 0.616 | 0.242 | 0.076 | 1.060 | 1.030 | 209 | - | - |
| $\beta_{ftmed2m}$ | Population recruitment | 0.018 | 0.052 | -0.041 | 0.161 | 1.000 | 7019 | 0.532 | 0.499 |
| $\beta_{fRHmax}$ | Population recruitment | -0.001 | 0.035 | -0.075 | 0.072 | 1.000 | 33757 | 0.487 | 0.500 |
| $\beta_{fsol}$ | Population recruitment | -0.006 | 0.046 | -0.105 | 0.065 | 1.020 | 15397 | 0.500 | 0.500 |
| $\beta_{ftmed0cm}$ | Population recruitment | 0.007 | 0.038 | -0.058 | 0.104 | 1.010 | 11688 | 0.499 | 0.500 |
| $\beta_{ftmin0cm}$ | Population recruitment | 0.008 | 0.044 | -0.054 | 0.108 | 1.010 | 15768 | 0.501 | 0.500 |
| $\beta_{fprecip}$ | Population recruitment | 0.001 | 0.038 | -0.073 | 0.079 | 1.010 | 32064 | 0.490 | 0.500 |
| $\beta_{fperf}$ | Population recruitment | 0.005 | 0.038 | -0.061 | 0.094 | 1.000 | 18827 | 0.495 | 0.500 |
| $\beta_{fha}$ | Population recruitment | 0.000 | 0.036 | -0.076 | 0.077 | 1.000 | 21077 | 0.493 | 0.500 |
| $\beta_{ffire}$ | Population recruitment | -0.001 | 0.043 | -0.089 | 0.078 | 1.000 | 119942 | 0.499 | 0.500 |
| $\beta_{fTSLF}$ | Population recruitment | -0.014 | 0.028 | -0.090 | 0.023 | 1.000 | 7174 | 0.512 | 0.500 |
| $\alpha_{p(C)}$ | Population capture | -3.541 | 0.191 | -3.918 | -3.167 | 1.000 | 3567 | - | - |

| Parameter | Vital rate | Mean | SD | 2.50% | 97.50% | $\widehat{R}$ | Effective sample size | Mean importance | SD importance |
|---|---|---|---|---|---|---|---|---|---|
| $\alpha_{p(Q)}$ | Population capture | -3.071 | 0.147 | -3.362 | -2.788 | 1.000 | 6560 | - | - |
| $\alpha_{p(EB)}$ | Population capture | -3.259 | 0.181 | -3.621 | -2.910 | 1.000 | 4713 | - | - |
| $\alpha_{p(MB)}$ | Population capture | -4.124 | 0.320 | -4.772 | -3.520 | 1.000 | 2617 | - | - |
| $\alpha_{p(LB)}$ | Population capture | -3.235 | 0.162 | -3.558 | -2.922 | 1.000 | 7213 | - | - |
| $\epsilon_{p(plot,t)}$ | Population capture | 0.494 | 0.050 | 0.395 | 0.591 | 1.000 | 5806 | - | - |
| **$\beta_{ptmed2m}$** | **Population capture** | **0.118** | **0.151** | **-0.033** | **0.475** | **1.000** | **1582** | **0.666** | **0.472** |
| **$\beta_{pRHmax}$** | **Population capture** | **0.128** | **0.144** | **-0.028** | **0.452** | **1.000** | **1851** | **0.714** | **0.452** |
| **$\beta_{psol}$** | **Population capture** | **0.055** | **0.065** | **-0.008** | **0.203** | **1.000** | **12028** | **0.619** | **0.486** |
| $\beta_{ptmed0cm}$ | Population capture | -0.006 | 0.074 | -0.211 | 0.132 | 1.010 | 5099 | 0.413 | 0.492 |
| $\beta_{ptmin0cm}$ | Population capture | -0.063 | 0.123 | -0.402 | 0.080 | 1.000 | 3120 | 0.568 | 0.495 |
| **$\beta_{pprecip}$** | **Population capture** | **-0.065** | **0.070** | **-0.210** | **0.011** | **1.000** | **5696** | **0.647** | **0.478** |
| **$\beta_{pperf}$** | **Population capture** | **-0.204** | **0.207** | **-0.691** | **0.002** | **1.000** | **1476** | **0.825** | **0.380** |
| $\beta_{pha}$ | Population capture | 0.038 | 0.048 | -0.006 | 0.147 | 1.000 | 18681 | 0.559 | 0.497 |

| Parameter | Vital rate | Mean | SD | 2.50% | 97.50% | $\widehat{R}$ | Effective sample size | Mean importance | SD importance |
|---|---|---|---|---|---|---|---|---|---|
| $\beta_{pfire}$ | Population capture | 0.054 | 0.146 | -0.124 | 0.488 | 1.000 | 15503 | 0.525 | 0.499 |
| $\beta_{pTSLF}$ | Population capture | -0.029 | 0.053 | -0.170 | 0.030 | 1.000 | 20646 | 0.451 | 0.498 |
| ***Micrablepharus atticolus*** | | | | | | | | | |
| $\alpha_{\sigma(C)}$ | Population survival | 2.873 | 0.526 | 1.919 | 3.970 | 1.110 | 208 | - | - |
| $\alpha_{\sigma(Q)}$ | Population survival | 2.645 | 0.326 | 2.091 | 3.342 | 1.070 | 279 | - | - |
| $\alpha_{\sigma(EB)}$ | Population survival | 2.128 | 0.306 | 1.584 | 2.749 | 1.060 | 214 | - | - |
| $\alpha_{\sigma(MB)}$ | Population survival | 2.291 | 0.271 | 1.817 | 2.877 | 1.080 | 265 | - | - |
| $\alpha_{\sigma(LB)}$ | Population survival | 1.369 | 0.331 | 0.742 | 2.059 | 1.090 | 142 | - | - |
| $\epsilon_{\sigma(plot,t)}$ | Population survival | 1.452 | 0.244 | 0.987 | 1.915 | 1.060 | 230 | - | - |
| $\beta_{\sigma tmed2m}$ | Population survival | 0.032 | 0.113 | -0.165 | 0.325 | 1.010 | 1191 | 0.503 | 0.500 |
| $\beta_{\sigma RHmax}$ | Population survival | -0.026 | 0.138 | -0.430 | 0.175 | 1.020 | 1538 | 0.489 | 0.500 |
| $\beta_{\sigma sol}$ | Population survival | 0.055 | 0.135 | -0.094 | 0.445 | 1.010 | 1052 | 0.515 | 0.500 |
| $\beta_{\sigma tmed0cm}$ | Population survival | 0.004 | 0.117 | -0.237 | 0.275 | 1.000 | 1994 | 0.466 | 0.499 |
| $\beta_{tmin0cm}$ | Population survival | 0.067 | 0.140 | -0.093 | 0.454 | 1.010 | 810 | 0.563 | 0.496 |

| Parameter | Vital rate | Mean | SD | 2.50% | 97.50% | $\hat{R}$ | Effective sample size | Mean importance | SD importance |
|---|---|---|---|---|---|---|---|---|---|
| **$\beta_{\sigma precip}$** | **Population survival** | **0.211** | **0.300** | **-0.029** | **1.005** | **1.050** | **285** | **0.736** | **0.441** |
| $\beta_{\sigma perf}$ | Population survival | 0.020 | 0.131 | -0.297 | 0.302 | 1.020 | 1158 | 0.518 | 0.500 |
| $\beta_{\sigma ha}$ | Population survival | 0.071 | 0.151 | -0.096 | 0.503 | 1.010 | 709 | 0.560 | 0.496 |
| $\beta_{\sigma fire}$ | Population survival | 0.029 | 0.185 | -0.241 | 0.496 | 1.030 | 6718 | 0.504 | 0.500 |
| $\beta_{\sigma TSLF}$ | Population survival | -0.004 | 0.050 | -0.133 | 0.100 | 1.000 | 2703 | 0.374 | 0.484 |
| $\alpha_{f(C)}$ | Population recruitment | -2.438 | 0.431 | -3.321 | -1.630 | 1.090 | 181 | - | - |
| $\alpha_{f(Q)}$ | Population recruitment | -2.282 | 0.227 | -2.815 | -1.908 | 1.080 | 184 | - | - |
| $\alpha_{f(EB)}$ | Population recruitment | -1.892 | 0.200 | -2.345 | -1.558 | 1.040 | 195 | - | - |
| $\alpha_{f(MB)}$ | Population recruitment | -2.045 | 0.211 | -2.514 | -1.696 | 1.030 | 162 | - | - |
| $\alpha_{f(LB)}$ | Population recruitment | -1.556 | 0.260 | -2.100 | -1.107 | 1.060 | 81 | - | - |
| $\epsilon_{f(plot,t)}$ | Population recruitment | 0.791 | 0.215 | 0.363 | 1.224 | 1.030 | 246 | - | - |
| $\beta_{ftmed2m}$ | Population recruitment | -0.002 | 0.209 | -0.628 | 0.377 | 1.080 | 146 | 0.441 | 0.496 |
| $\beta_{fRHmax}$ | Population recruitment | -0.106 | 0.252 | -0.819 | 0.237 | 1.060 | 408 | 0.491 | 0.500 |

| Parameter | Vital rate | Mean | SD | 2.50% | 97.50% | $\widehat{R}$ | Effective sample size | Mean importance | SD importance |
|---|---|---|---|---|---|---|---|---|---|
| $\beta_{fsol}$ | Population recruitment | 0.092 | 0.194 | -0.192 | 0.580 | 1.040 | 228 | 0.500 | 0.500 |
| **$\beta_{ftmed0cm}$** | **Population recruitment** | **0.501** | **0.436** | **0.000** | **1.475** | **1.090** | **58** | **0.825** | **0.380** |
| **$\beta_{ftmin0cm}$** | **Population recruitment** | **0.215** | **0.339** | **-0.118** | **1.169** | **1.070** | **182** | **0.593** | **0.491** |
| **$\beta_{fprecip}$** | **Population recruitment** | **0.189** | **0.262** | **-0.075** | **0.823** | **1.020** | **505** | **0.591** | **0.492** |
| **$\beta_{fperf}$** | **Population recruitment** | **-0.230** | **0.500** | **-1.703** | **0.307** | **1.050** | **188** | **0.553** | **0.497** |
| **$\beta_{fha}$** | **Population recruitment** | **0.439** | **0.680** | **-0.137** | **2.339** | **1.030** | **119** | **0.650** | **0.477** |
| **$\beta_{ffire}$** | **Population recruitment** | **0.195** | **0.524** | **-0.474** | **1.755** | **1.010** | **1979** | **0.548** | **0.498** |
| $\beta_{fTSLF}$ | Population recruitment | 0.004 | 0.037 | -0.070 | 0.112 | 1.020 | 1659 | 0.215 | 0.411 |
| $\alpha_{p(C)}$ | Population capture | -4.568 | 0.494 | -5.540 | -3.607 | 1.030 | 440 | - | - |
| $\alpha_{p(Q)}$ | Population capture | -3.361 | 0.174 | -3.704 | -3.024 | 1.010 | 1194 | - | - |
| $\alpha_{p(EB)}$ | Population capture | -3.131 | 0.190 | -3.507 | -2.763 | 1.010 | 1154 | - | - |
| $\alpha_{p(MB)}$ | Population capture | -2.820 | 0.154 | -3.120 | -2.517 | 1.000 | 1018 | - | - |
| $\alpha_{p(LB)}$ | Population capture | -3.360 | 0.280 | -3.912 | -2.814 | 1.010 | 780 | - | - |

| Parameter | Vital rate | Mean | SD | 2.50% | 97.50% | $\widehat{R}$ | Effective sample size | Mean importance | SD importance |
|---|---|---|---|---|---|---|---|---|---|
| $\epsilon_{p(plot,t)}$ | Population capture | 0.539 | 0.064 | 0.416 | 0.665 | 1.000 | 3449 | - | - |
| $\beta_{ptmed2m}$ | Population capture | 0.025 | 0.121 | -0.212 | 0.357 | 1.000 | 3160 | 0.412 | 0.492 |
| $\beta_{pRHmax}$ | Population capture | -0.008 | 0.094 | -0.253 | 0.204 | 1.000 | 7002 | 0.368 | 0.482 |
| $\beta_{psol}$ | Population capture | -0.022 | 0.059 | -0.194 | 0.051 | 1.000 | 10287 | 0.312 | 0.463 |
| $\boldsymbol{\beta_{ptmed0cm}}$ | **Population capture** | **-0.248** | **0.166** | **-0.589** | **0.000** | **1.000** | **2991** | **0.859** | **0.348** |
| $\boldsymbol{\beta_{ptmin0cm}}$ | **Population capture** | **-0.357** | **0.193** | **-0.773** | **0.000** | **1.010** | **2344** | **0.933** | **0.249** |
| $\boldsymbol{\beta_{pprecip}}$ | **Population capture** | **-0.415** | **0.106** | **-0.626** | **-0.211** | **1.010** | **2334** | **0.999** | **0.032** |
| $\beta_{pperf}$ | Population capture | 0.074 | 0.156 | -0.138 | 0.491 | 1.010 | 2940 | 0.476 | 0.499 |
| $\beta_{pha}$ | Population capture | -0.065 | 0.128 | -0.402 | 0.103 | 1.000 | 4912 | 0.459 | 0.498 |
| $\beta_{pfire}$ | Population capture | 0.070 | 0.181 | -0.224 | 0.549 | 1.000 | 38116 | 0.490 | 0.500 |
| $\boldsymbol{\beta_{pTSLF}}$ | **Population capture** | **-0.097** | **0.118** | **-0.358** | **0.017** | **1.000** | **3976** | **0.576** | **0.494** |
| | | | | ***Tropidurus itambere*** | | | | | |
| $\alpha_{\sigma(C)}$ | Population survival | 1.670 | 0.451 | 0.849 | 2.616 | 1.020 | 411 | - | - |
| $\alpha_{\sigma(Q)}$ | Population survival | 1.910 | 0.168 | 1.604 | 2.264 | 1.040 | 598 | - | - |

| Parameter | Vital rate | Mean | SD | 2.50% | 97.50% | $\widehat{R}$ | Effective sample size | Mean importance | SD importance |
|---|---|---|---|---|---|---|---|---|---|
| $\alpha_{\sigma(EB)}$ | Population survival | 1.521 | 0.149 | 1.256 | 1.853 | 1.020 | 363 | - | - |
| $\alpha_{\sigma(MB)}$ | Population survival | 1.268 | 0.143 | 1.008 | 1.561 | 1.030 | 272 | - | - |
| $\alpha_{\sigma(LB)}$ | Population survival | 1.430 | 0.158 | 1.156 | 1.769 | 1.020 | 276 | - | - |
| $\epsilon_{\sigma(plot,t)}$ | Population survival | 0.396 | 0.218 | 0.051 | 0.847 | 1.100 | 133 | - | - |
| **$\beta_{\sigma tmed2m}$** | **Population survival** | **-0.165** | **0.262** | **-0.835** | **0.059** | **1.040** | **389** | **0.633** | **0.482** |
| $\beta_{\sigma RHmax}$ | Population survival | 0.065 | 0.157 | -0.076 | 0.538 | 1.010 | 774 | 0.519 | 0.500 |
| **$\beta_{\sigma sol}$** | **Population survival** | **0.262** | **0.333** | **-0.028** | **1.033** | **1.030** | **308** | **0.730** | **0.444** |
| $\beta_{\sigma tmed0cm}$ | Population survival | -0.027 | 0.115 | -0.365 | 0.126 | 1.020 | 1124 | 0.453 | 0.498 |
| $\beta_{tmin0cm}$ | Population survival | -0.029 | 0.140 | -0.490 | 0.146 | 1.010 | 569 | 0.475 | 0.499 |
| **$\beta_{\sigma precip}$** | **Population survival** | **0.242** | **0.272** | **-0.014** | **0.864** | **1.040** | **253** | **0.790** | **0.408** |
| $\beta_{\sigma perf}$ | Population survival | -0.006 | 0.133 | -0.338 | 0.204 | 1.040 | 731 | 0.462 | 0.499 |
| $\beta_{\sigma ha}$ | Population survival | -0.007 | 0.092 | -0.262 | 0.149 | 1.000 | 1403 | 0.431 | 0.495 |
| $\beta_{\sigma fire}$ | Population survival | -0.019 | 0.200 | -0.567 | 0.349 | 1.010 | 20641 | 0.497 | 0.500 |

| **Parameter** | **Vital rate** | **Mean** | **SD** | **2.50%** | **97.50%** | $\hat{R}$ | **Effective sample size** | **Mean importance** | **SD importance** |
|---|---|---|---|---|---|---|---|---|---|
| $\beta_{\sigma TSLF}$ | Population survival | -0.002 | 0.028 | -0.074 | 0.056 | 1.000 | 4803 | 0.318 | 0.466 |
| $\alpha_{f(C)}$ | Population recruitment | -2.182 | 0.520 | -3.410 | -1.360 | 1.020 | 278 | - | - |
| $\alpha_{f(Q)}$ | Population recruitment | -2.344 | 0.252 | -2.887 | -1.918 | 1.060 | 201 | - | - |
| $\alpha_{f(EB)}$ | Population recruitment | -2.019 | 0.231 | -2.516 | -1.625 | 1.040 | 108 | - | - |
| $\alpha_{f(MB)}$ | Population recruitment | -1.816 | 0.205 | -2.270 | -1.478 | 1.050 | 90 | - | - |
| $\alpha_{f(LB)}$ | Population recruitment | -1.969 | 0.237 | -2.521 | -1.568 | 1.020 | 88 | - | - |
| $\epsilon_{f(plot,t)}$ | Population recruitment | 1.085 | 0.250 | 0.638 | 1.612 | 1.050 | 195 | - | - |
| $\boldsymbol{\beta_{ftmed2m}}$ | **Population recruitment** | **0.437** | **0.225** | **0.011** | **0.896** | **1.010** | **536** | 1* | 0* |
| $\beta_{fRHmax}$ | Population recruitment | 0.000 | 0.000 | 0.000 | 0.000 | NA | 0 | 0* | 0* |
| $\boldsymbol{\beta_{fsol}}$ | **Population recruitment** | **-0.218** | **0.290** | **-0.801** | **0.344** | **1.010** | **600** | 1* | 0* |
| $\beta_{ftmed0cm}$ | Population recruitment | 0.000 | 0.000 | 0.000 | 0.000 | NA | 0 | 0* | 0* |
| $\beta_{ftmin0cm}$ | Population recruitment | 0.000 | 0.000 | 0.000 | 0.000 | NA | 0 | 0* | 0* |
| $\beta_{fprecip}$ | Population recruitment | 0.000 | 0.000 | 0.000 | 0.000 | NA | 0 | 0* | 0* |

| Parameter | Vital rate | Mean | SD | 2.50% | 97.50% | $\widehat{R}$ | Effective sample size | Mean importance | SD importance |
|---|---|---|---|---|---|---|---|---|---|
| $\beta_{fperf}$ | Population recruitment | 0.000 | 0.000 | 0.000 | 0.000 | NA | 0 | 0* | 0* |
| $\beta_{fha}$ | Population recruitment | 0.000 | 0.000 | 0.000 | 0.000 | NA | 0 | 0* | 0* |
| $\beta_{ffire}$ | Population recruitment | 0.000 | 0.000 | 0.000 | 0.000 | NA | 0 | 0* | 0* |
| $\beta_{fTSLF}$ | Population recruitment | 0.000 | 0.000 | 0.000 | 0.000 | NA | 0 | 0* | 0* |
| $\alpha_{p(C)}$ | Population capture | -4.435 | 0.625 | -5.735 | -3.285 | 1.000 | 2704 | - | - |
| $\alpha_{p(Q)}$ | Population capture | -3.245 | 0.176 | -3.597 | -2.907 | 1.010 | 4541 | - | - |
| $\alpha_{p(EB)}$ | Population capture | -2.854 | 0.159 | -3.170 | -2.545 | 1.000 | 3633 | - | - |
| $\alpha_{p(MB)}$ | Population capture | -2.490 | 0.150 | -2.789 | -2.202 | 1.010 | 4049 | - | - |
| $\alpha_{p(LB)}$ | Population capture | -2.779 | 0.169 | -3.118 | -2.451 | 1.000 | 4239 | - | - |
| $\epsilon_{p(plot,t)}$ | Population capture | 0.512 | 0.058 | 0.396 | 0.622 | 1.010 | 2996 | - | - |
| **$\beta_{ptmed2m}$** | **Population capture** | **0.130** | **0.140** | **-0.015** | **0.440** | **1.000** | **2468** | **0.646** | **0.478** |
| $\beta_{pRHmax}$ | Population capture | -0.037 | 0.087 | -0.279 | 0.077 | 1.000 | 5174 | 0.371 | 0.483 |
| **$\beta_{psol}$** | **Population capture** | **-0.264** | **0.088** | **-0.438** | **-0.090** | **1.000** | **4381** | **0.990** | **0.100** |

| Parameter | Vital rate | Mean | SD | 2.50% | 97.50% | $\widehat{R}$ | Effective sample size | Mean importance | SD importance |
|---|---|---|---|---|---|---|---|---|---|
| $\boldsymbol{\beta_{ptmed0cm}}$ | **Population capture** | **0.145** | **0.189** | **-0.046** | **0.622** | **1.000** | **1168** | **0.601** | **0.490** |
| $\boldsymbol{\beta_{ptmin0cm}}$ | **Population capture** | **-0.158** | **0.249** | **-0.814** | **0.124** | **1.010** | **923** | **0.582** | **0.493** |
| $\boldsymbol{\beta_{pprecip}}$ | **Population capture** | **-0.500** | **0.097** | **-0.698** | **-0.316** | **1.000** | **3677** | **1.000** | **0.000** |
| $\boldsymbol{\beta_{pperf}}$ | **Population capture** | **0.214** | **0.296** | **-0.069** | **1.015** | **1.010** | **702** | **0.631** | **0.483** |
| $\beta_{pha}$ | Population capture | 0.008 | 0.048 | -0.084 | 0.147 | 1.000 | 10852 | 0.254 | 0.435 |
| $\boldsymbol{\beta_{pfire}}$ | **Population capture** | **-0.173** | **0.243** | **-0.775** | **0.112** | **1.000** | **37961** | **0.611** | **0.487** |
| $\beta_{pTSLF}$ | Population capture | -0.007 | 0.070 | -0.196 | 0.152 | 1.000 | 23746 | 0.308 | 0.462 |

* Fixed values

As mentioned in the main text, the environmental variables were poor predictors of the *C. nigropunctatum* vital rates. In *M. atticolus*, precipitation affected positively survival (Table S4). *T. itambere* survival decreased with high mean temperatures (*tmed2m*) and increased with high insolation and precipitation (Table S4). Recruitment of *M. atticolus* decreased in months with higher mean locomotor performance and increased in months/plots with fire and high temperatures (*tmed2m* and *tmin0cm*) and hours of activity. Recruitment in *T. itambere* increased with high mean temperatures (*tmed2m*) and low insolation (Table S4). Although most vital rates were not affected by microclimatic or ecophysiological variables, capture probabilities were in all species. In *C. nigropunctatum*, capture probability was positively affected by mean air temperature, air humidity, and insolation and negatively affected by precipitation and mean locomotor performance (*tmed2m*, *RHmax*, *sol*, *precip*, and *perf;* Table S4). In *M. atticolus*, capture probabilities decreased with high temperatures and precipitation and with time since the last fire (*tmed0cm*, *tmin0cm*, *precip*, *TSLF*; Table S4). In *T. itambere*, capture probability increased in months/plots with high mean temperatures (*tmed2m* and *tmed0cm*) and mean locomotor performance (*perf*) and decreased with high minimum temperatures, insolation, and precipitation (*tmin0cm*, *sol*, and *precip*; Table S4). The capture probability of *T. itambere* also decreased in months when fires occurred (Table S4).

**Survival and capture probabilities and Goodness of fit tests**

Here, we present the results of the mean estimates of populations' survival and captures from the three lizard species among the studied plots. Considering our time resolution (months) and credibility levels, we have high confidence in the estimates of the vital rates our models provide.

**Table S5.** Model-averaged means and credible intervals (2.5% and 97.5%) of populations' survival ($\sigma$) and capture ($p$) of three lizard species from the Brazilian Cerrado savannas (*Copeoglossum nigropunctatum*, *Micrablepharus atticolus* and *Tropidurus itambere*). Populations were monitored in five plots submitted to fire regimes of varying severity: Control (C), Quadrennial (Q), Early Biennial (EB), Mid Biennial (MB), and Late Biennial (LB).

| **Parameter** | **Mean** | **2.50%** | **97.50%** |
|---|---|---|---|
| | *C. nigropunctatum* | | |
| $\sigma$(C) | 0.918 | 0.879 | 0.953 |
| $\sigma$(Q) | 0.918 | 0.881 | 0.951 |
| $\sigma$(EB) | 0.908 | 0.864 | 0.946 |
| $\sigma$(MB) | 0.925 | 0.870 | 0.962 |
| $\sigma$(LB) | 0.930 | 0.895 | 0.959 |
| $p$(C) | 0.028 | 0.019 | 0.040 |
| $p$(Q) | 0.044 | 0.034 | 0.058 |
| $p$(EB) | 0.037 | 0.026 | 0.052 |
| $p$(MB) | 0.016 | 0.008 | 0.029 |
| $p$(LB) | 0.038 | 0.028 | 0.051 |

| Parameter | Mean | 2.50% | 97.50% |
|---|---|---|---|
| *M. atticolus* | | | |
| $\sigma$(C) | 0.946 | 0.872 | 0.981 |
| $\sigma$(Q) | 0.934 | 0.890 | 0.966 |
| $\sigma$(EB) | 0.894 | 0.830 | 0.940 |
| $\sigma$(MB) | 0.908 | 0.860 | 0.947 |
| $\sigma$(LB) | 0.797 | 0.677 | 0.887 |
| $p$(C) | 0.010 | 0.004 | 0.026 |
| $p$(Q) | 0.034 | 0.024 | 0.046 |
| $p$(EB) | 0.042 | 0.029 | 0.059 |
| $p$(MB) | 0.056 | 0.042 | 0.075 |
| $p$(LB) | 0.034 | 0.020 | 0.057 |
| *T. itambere* | | | |
| $\sigma$(C) | 0.842 | 0.700 | 0.932 |
| $\sigma$(Q) | 0.871 | 0.833 | 0.906 |
| $\sigma$(EB) | 0.821 | 0.778 | 0.864 |
| $\sigma$(MB) | 0.780 | 0.733 | 0.827 |
| $\sigma$(LB) | 0.807 | 0.761 | 0.854 |
| $p$(C) | 0.012 | 0.003 | 0.036 |
| $p$(Q) | 0.038 | 0.027 | 0.052 |
| $p$(EB) | 0.054 | 0.040 | 0.073 |
| $p$(MB) | 0.077 | 0.058 | 0.100 |
| $p$(LB) | 0.058 | 0.042 | 0.079 |

To date, no goodness-of-fit (GOF) tests for models with individual time-varying covariates or temporal covariates, as the ones we used here, exist. Equally, GOF tests for PJS models do not exist that we are aware of. Indeed, because our models allowed for survival and capture probabilities to vary by age and size per individual, they can control for some issues of heterogeneity in the capture history, such as trap dependence and transient individuals. However, we present here the results of the GOF tests for CJS models using the package R2UCARE (Choquet *et al.* 2009; Gimenez *et al.* 2018) to demonstrate that even simpler models do not compromise the model's assumptions of homogeneity in survival and re(capture) probability of the individuals (Table S6). The overall GOF tests whether the CJS model adequately fits the data using resampling methods and deviance as a metric (White 2002). Test 3.SR tests whether newly encountered individuals have the same chance to be later reobserved as recaptured (previously encountered) individuals, *i.e.*, tests for transient individuals; test 2.CT tests whether missed individuals have the same chance to be recaptured on the next occasion as currently captured individuals, *i.e.*, tests for trap-dependence (trap-shyness or trap-happiness, *sensu* (Pradel 1993); test 3.SM tests whether, among those individuals seen again, the occasion when they were seen does not differ among previously and newly marked individuals, *i.e.*, tests for differences in the time for the recapture between newly and previously marked individuals; and the test 2.CL tests whether there is a difference in the timing of reencounters between the individuals encountered and not encountered at occasion $i$, conditional on presence at both occasions $i$ and $i + 2$ (Pollock, Hines & Nichols 1985; Pradel 1993; Gimenez *et al.* 2018). We only found significant differences for the test for trap-dependence (test 2.CT) for *T. itambere*, considering all the plots pooled together (Table S6). The sign of the statistic was negative (-5.593), indicating that missed individuals have a lower chance of being recaptured on the next occasion than currently captured individuals (*i.e.*, trap-happiness *sensu* (Pradel 1993). However, analyzing each plot separately, we found no significantly different probabilities of missed individuals being recaptured on the next occasion than currently captured individuals (Table S6). Notice that in all GOF tests in our three species, we have a higher number of degrees of freedom when compared to the statistic (in this case, $\chi^2$), indicating underdispersion in our data. The underdispersion likely happens because of our high temporal resolution (months), which decreases the variance concerning the mean of capture probability. This kind of data may generate overfitted models (Sellers & Morris 2017). Still, we built our Bayesian and hierarchical approach to address these potential issues, such as incorporating individual covariates (size) and temporal random variation (see the section on models' parameterizations).

**Table S6.** Goodness-of-fit (GOF) tests for the Cormack-Jolly-Seber Models (CJS) for three species of lizards from the Brazilian Cerrado savannas (*Copeoglossum nigropunctatum*, *Micrablepharus atticolus*, and *Tropidurus itambere*) in five plots submitted to fire regimes of varying severity: Control (C), Quadrennial (Q), Early Biennial (EB), Mid Biennial (MB), and Late Biennial (LB).

| Plot | Test | Statistic | DF | *P* |
|---|---|---|---|---|
| | *C. nigropunctatum* | | | |
| All | Overall GOF test | 186.806 | 369 | 1 |
| C | Overall GOF test | 18.983 | 80 | 1 |
| Q | Overall GOF test | 41.253 | 136 | 1 |
| EB | Overall GOF test | 24.207 | 85 | 1 |
| MB | Overall GOF test | 1.644 | 22 | 1 |
| LB | Overall GOF test | 34.589 | 118 | 1 |
| All | Test 3.SR - transients | 53.171 | 102 | 1 |
| C | Test 3.SR - transients | 9.417 | 22 | 0.991 |
| Q | Test 3.SR - transients | 7.321 | 42 | 1 |
| EB | Test 3.SR - transients | 8.125 | 21 | 0.995 |
| MB | Test 3.SR - transients | 0 | 3 | 1 |
| LB | Test 3.SR - transients | 9.633 | 33 | 1 |
| All | Test 2.CT - trap-dependence | 59.035 | 102 | 1 |
| C | Test 2.CT - trap-dependence | 3.791 | 21 | 1 |
| Q | Test 2.CT - trap-dependence | 16.598 | 39 | 0.999 |
| EB | Test 2.CT - trap-dependence | 8.57 | 24 | 0.998 |
| MB | Test 2.CT - trap-dependence | 0.936 | 7 | 0.996 |
| LB | Test 2.CT - trap-dependence | 16.874 | 32 | 0.987 |
| All | Test 3.SM - long-term transients | 7.949 | 48 | 1 |
| C | Test 3.SM - long-term transients | 0 | 3 | 1 |
| Q | Test 3.SM - long-term transients | 0.936 | 7 | 0.996 |
| EB | Test 3.SM - long-term transients | 0 | 2 | 1 |
| MB | Test 3.SM - long-term transients | 0 | 1 | 1 |
| LB | Test 3.SM - long-term transients | 0 | 6 | 1 |
| All | Test 2.CL - long-term trap-dependence | 66.651 | 117 | 1 |
| C | Test 2.CL - long-term trap-dependence | 5.775 | 34 | 1 |
| Q | Test 2.CL - long-term trap-dependence | 16.398 | 48 | 1 |
| EB | Test 2.CL - long-term trap-dependence | 7.512 | 38 | 1 |
| MB | Test 2.CL - long-term trap-dependence | 0.708 | 11 | 1 |
| LB | Test 2.CL - long-term trap-dependence | 8.082 | 47 | 1 |
| | *M. atticolus* | | | |
| All | Overall GOF test | 115.251 | 188 | 1 |
| C | Overall GOF test | 0 | 3 | 1 |
| Q | Overall GOF test | 12.727 | 44 | 1 |

| Plot | Test | Statistic | DF | *P* |
|---|---|---|---|---|
| EB | Overall GOF test | 9.076 | 42 | 1 |
| MB | Overall GOF test | 31.894 | 96 | 1 |
| LB | Overall GOF test | 4.406 | 11 | 0.957 |
| All | Test 3.SR - transients | 31.069 | 56 | 0.997 |
| C | Test 3.SR - transients | 0 | 1 | 1 |
| Q | Test 3.SR - transients | 5.23 | 13 | 0.97 |
| EB | Test 3.SR - transients | 4.558 | 15 | 0.995 |
| MB | Test 3.SR - transients | 8.487 | 33 | 1 |
| LB | Test 3.SR - transients | 4.406 | 6 | 0.622 |
| All | Test 2.CT - trap-dependence | 43.592 | 51 | 0.76 |
| C | Test 2.CT - trap-dependence | 0 | 2 | 1 |
| Q | Test 2.CT - trap-dependence | 5.853 | 15 | 0.982 |
| EB | Test 2.CT - trap-dependence | 1.323 | 10 | 0.999 |
| MB | Test 2.CT - trap-dependence | 17.282 | 27 | 0.924 |
| LB | Test 2.CT - trap-dependence | 0 | 4 | 1 |
| All | Test 3.SM - long-term transients | 13.585 | 22 | 0.916 |
| C | Test 3.SM - long-term transients | 0 | 0 | 1 |
| Q | Test 3.SM - long-term transients | 0.936 | 2 | 0.626 |
| EB | Test 3.SM - long-term transients | 0.936 | 5 | 0.968 |
| MB | Test 3.SM - long-term transients | 2.66 | 10 | 0.988 |
| LB | Test 3.SM - long-term transients | 0 | 0 | 1 |
| All | Test 2.CL - long-term trap-dependence | 27.005 | 59 | 1 |
| C | Test 2.CL - long-term trap-dependence | 0 | 0 | 1 |
| Q | Test 2.CL - long-term trap-dependence | 0.708 | 14 | 1 |
| EB | Test 2.CL - long-term trap-dependence | 2.259 | 12 | 0.999 |
| MB | Test 2.CL - long-term trap-dependence | 3.465 | 26 | 1 |
| LB | Test 2.CL - long-term trap-dependence | 0 | 1 | 1 |
| | *T. itambere* | | | |
| All | Overall GOF test | 249.575 | 347 | 1 |
| C | Overall GOF test | 0 | 0 | 1 |
| Q | Overall GOF test | 27.736 | 97 | 1 |
| EB | Overall GOF test | 39.881 | 113 | 1 |
| MB | Overall GOF test | 25.323 | 104 | 1 |
| LB | Overall GOF test | 38.607 | 93 | 1 |
| All | Test 3.SR - transients | 57.624 | 104 | 1 |
| C | Test 3.SR - transients | 0 | 0 | 1 |
| Q | Test 3.SR - transients | 8.7 | 33 | 1 |
| EB | Test 3.SR - transients | 9.974 | 40 | 1 |
| MB | Test 3.SR - transients | 15.44 | 38 | 1 |
| LB | Test 3.SR - transients | 10.643 | 29 | 0.999 |
| **All** | **Test 2.CT - trap-dependence** | **145.012** | **102** | **0.003** |
| C | Test 2.CT - trap-dependence | 0 | 0 | 1 |

| Plot | Test | Statistic | DF | *P* |
|---|---|---|---|---|
| Q | Test 2.CT - trap-dependence | 16.684 | 31 | 0.983 |
| EB | Test 2.CT - trap-dependence | 22.756 | 36 | 0.958 |
| MB | Test 2.CT - trap-dependence | 7.852 | 35 | 1 |
| LB | Test 2.CT - trap-dependence | 16.642 | 33 | 0.992 |
| All | Test 3.SM - long-term transients | 10.672 | 41 | 1 |
| C | Test 3.SM - long-term transients | 0 | 0 | 1 |
| Q | Test 3.SM - long-term transients | 0 | 5 | 1 |
| EB | Test 3.SM - long-term transients | 0 | 7 | 1 |
| MB | Test 3.SM - long-term transients | 0 | 6 | 1 |
| LB | Test 3.SM - long-term transients | 0.936 | 5 | 0.968 |
| All | Test 2.CL - long-term trap-dependence | 36.267 | 100 | 1 |
| C | Test 2.CL - long-term trap-dependence | 0 | 0 | 1 |
| Q | Test 2.CL - long-term trap-dependence | 2.352 | 28 | 1 |
| EB | Test 2.CL - long-term trap-dependence | 7.151 | 30 | 1 |
| MB | Test 2.CL - long-term trap-dependence | 2.031 | 25 | 1 |
| LB | Test 2.CL - long-term trap-dependence | 10.386 | 26 | 0.997 |

**Population growth**

We provide the population growth rates ($\lambda$) estimated from our IPMs and the PJS models across the 14 years of our study. As expected, they are highly correlated (Spearman correlation > 0.99 for all populations in all plots (Fig. S10). The realized population growth rates of *M. atticolus* and *T. itambere* have higher variance in the most severe fire regime (Fig. S11), which agrees that populations have lower resilience in this fire regime. *C. nigropunctatum* had higher variance (S.D.) in the intermediate fire regime (third level – early biennial fires; Fig. S11), where populations displayed lower resistance but higher compensation and faster recovery. As expected, the realized population growth rates (derived from the PJS models) show the same relationships as the reproductive output and demographic resilience components (Fig. S12). Resistance and recovery time decreased with increasing realized population growth rates, while compensation increased (Fig. S12). *M. atticolus* showed opposite relationships, with higher resistance and lower compensation when realized population growth rates increased (Fig. S12).

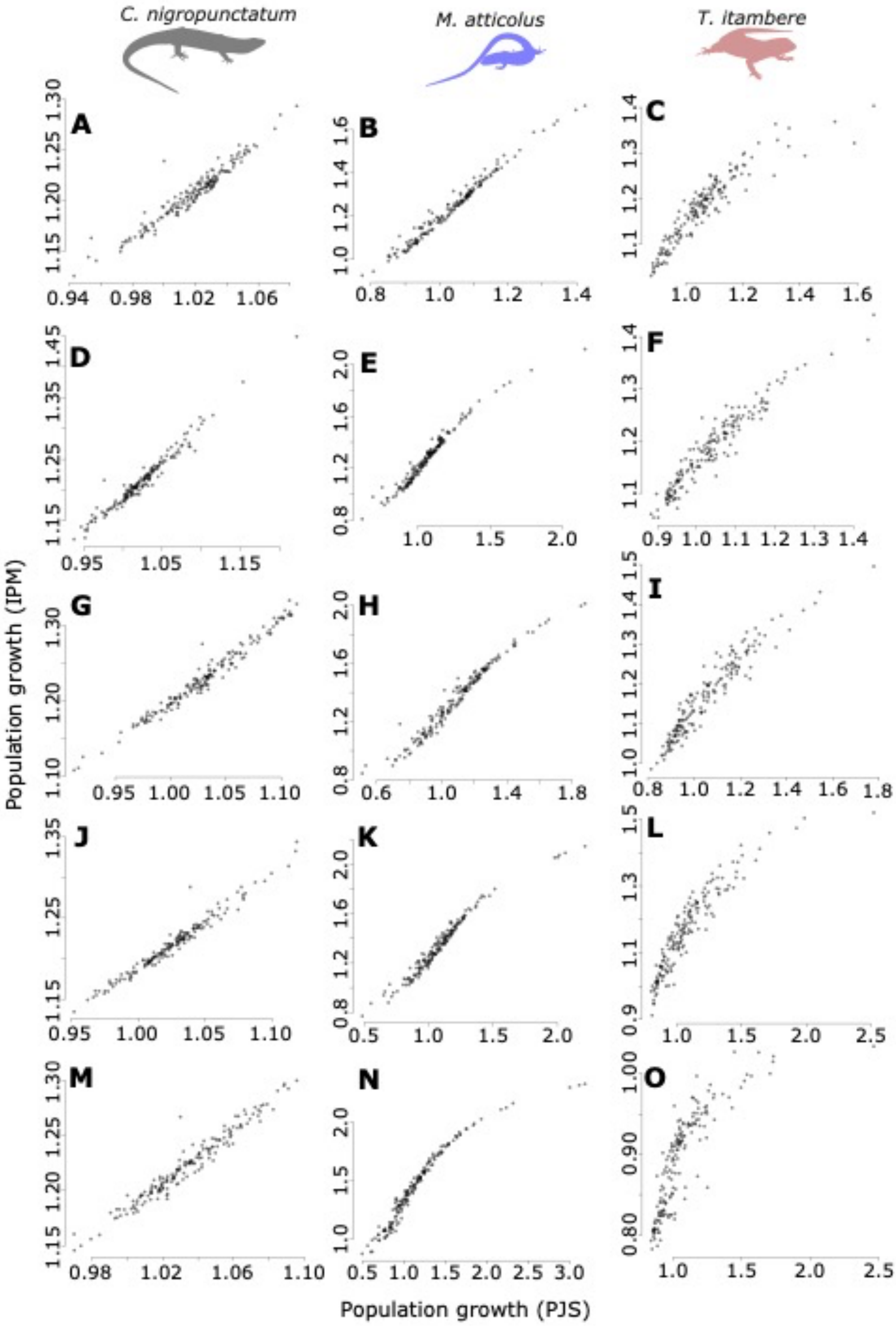


**Fig. S10.** Variation of population growth ($\lambda$) of three lizard species (*Copeoglossum nigropunctatum* – A, D, G, J, and M; *Micrablepharus atticolus* – B, E, H, K, and N; and *Tropidurus itambere* – C, F, I, L, and O) estimated from our Integral Projection Models (IPMs) and Pradel Jolly-Seber (PJS) models from the Brazilian Cerrado savannas in fire regimes of varying severity. Fire severity increases from top (A-C) to bottom (M-O).

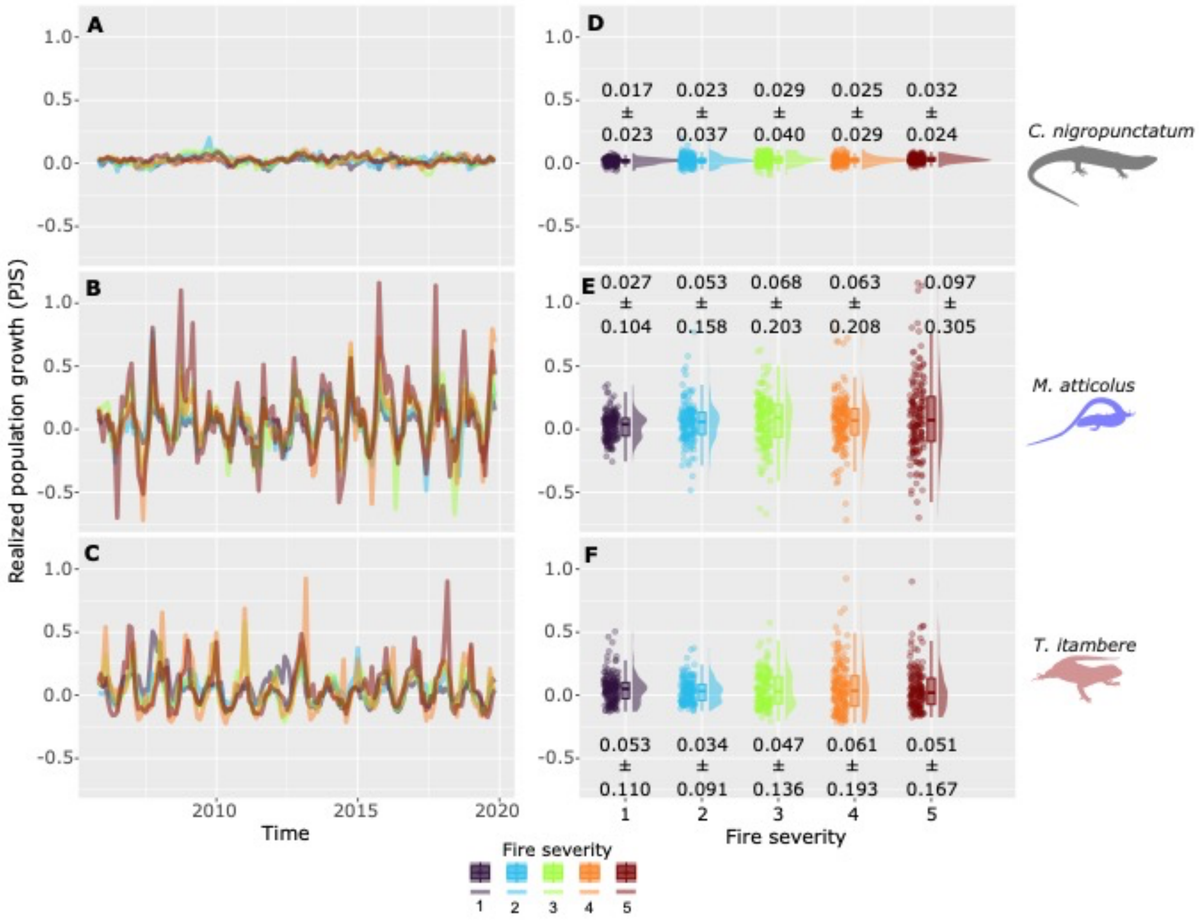


**Fig. S11.** Realized population growth rates in log-scale (*log(λ)*) from three species of lizards (*Copeoglossum nigropunctatum*, *Micrablepharus atticolus*, and *Tropidurus itambere*) over time (A, B, and C) and between fire regimes (D, E, and F) from the Brazilian Cerrado savannas. Using mark-recapture individual data, we derived population growth rates with Pradel Jolly-Seber (PJS) models. Boxplots depict the median (solid horizontal bars) and interquartile range (boxes). Whiskers extend as far as 1.5x the interquartile range or to minimum and maximum values. The vertical solid horizontal bars indicate the $90^{th}$ temperature percentiles. In D, E, and F, we show the mean ± S.D. for each population by species and plot (fire severity).

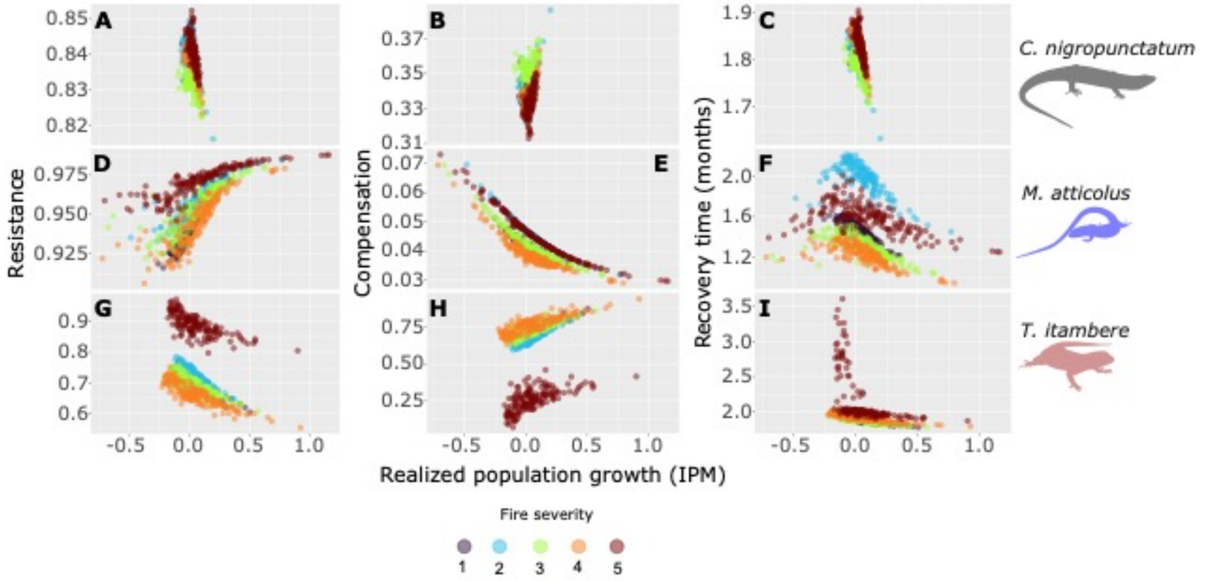


**Fig. S12.** Variation of realized population growth (*log(λ)*), estimated from Pradel Jolly-Seber (PJS) models) and demographic resilience components (resistance –A, D, and G; compensation – B, E, and H; and recovery time – C, F, and I) estimated from our Integral Projection Models (IPMs) of three lizard species (*Copeoglossum nigropunctatum* – A-C; *Micrablepharus atticolus* – D-F; and *Tropidurus itambere* – G-I) from the Brazilian Cerrado savannas in fire regimes of varying severity.

#### Life history traits

With the monthly stochastic Integral Projection Models, we estimated eight different life history traits of the studied species with package Rage(Jones *et al.* 2022): generation time; degree of iteroparity; life expectancy; variance of life expectancy; longevity; net reproductive output expected lifetime production of offspring that start life by an individual also starting life; net reproductive output calculated as the per-generation population growth rate; "shape" value of the distribution of reproduction over size; "shape" value of survival lifespan inequality; shape of the age-specific survivorship curve. The Principal Component Analysis (PCA) shows that species have different life history strategies, summarized by two main axes that account for 81.43% of the variation (Fig. S13). The first axis is related to the time schedules of mortality and reproduction, with populations with long-living individuals and reproduction more spread through life in one end and populations with short-living individuals and high reproductive output concentrated in much shorter time intervals (Fig. S13). The second axis is related to the fast-slow continuum, with populations with long generation time and low-reproductive output in one end and short generation time and high-reproductive output in another (Fig. S13).

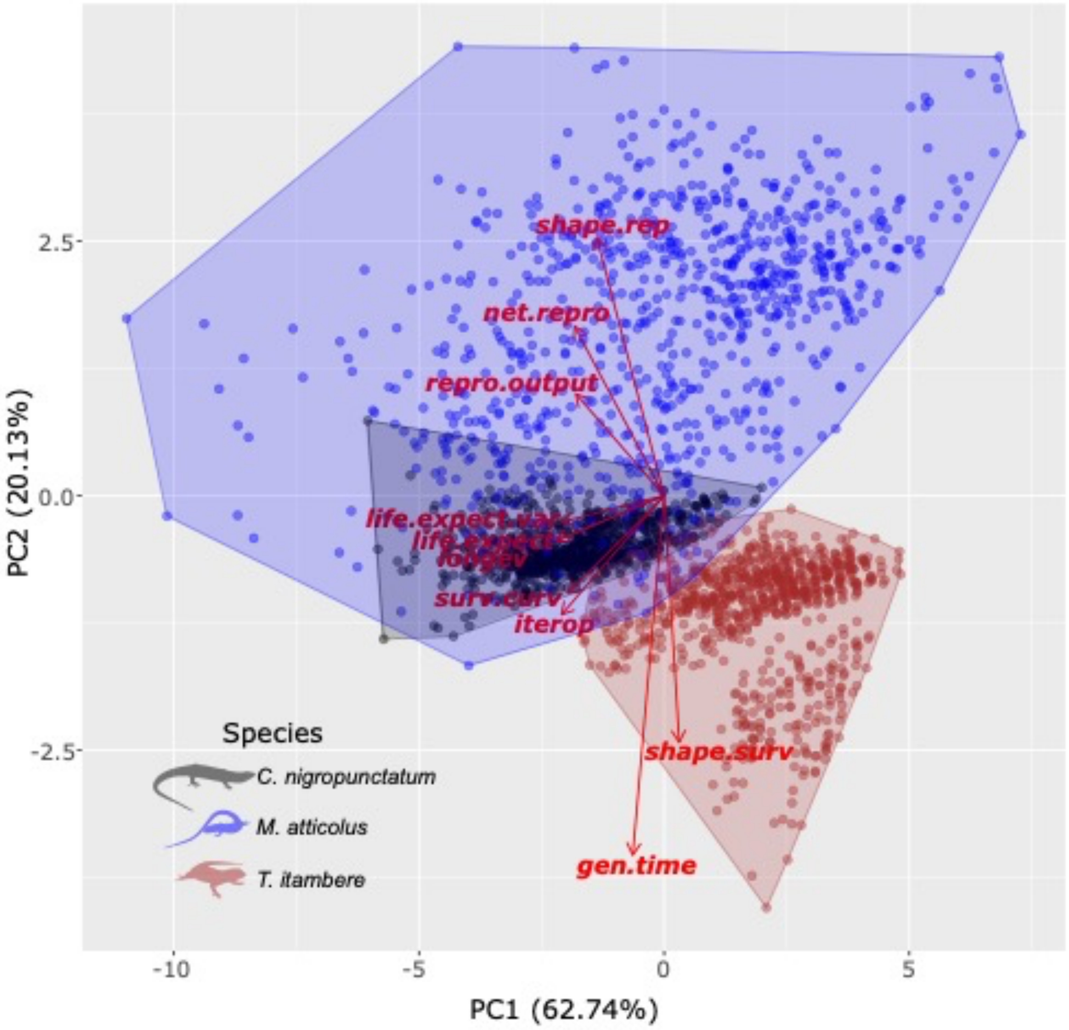


**Fig. S13.** Principal component analysis (PCA) showing the variation of life history traits from three species of lizards (*Copeoglossum nigropunctatum*, *Micrablepharus atticolus*, and *Tropidurus itambere*) from the Brazilian Cerrado savannas. *gen.time* = generation time; *iterop* = degree of iteroparity; *life.expect* = life expectancy; *life.expect.var* = variance of life expectancy; *longev* = longevity; *net.repro* = net reproductive output expected lifetime production of offspring that start life by an individual also starting life; *repro.output* = net reproductive output calculated as the per-generation population growth rate; *shape.rep* = 'shape' value of distribution of reproduction over size; *shape.surv* = 'shape' value of survival lifespan inequality; *surv.curv* = shape of the age-specific survivorship curve.

Life history traits are highly correlated (Fig. S14). For instance, life expectancy, longevity, and degree of iteroparity have more than 90% of positive correlation (Fig. S14). We also show that life history traits and demographic resilience components (resistance, compensation, damping ratio, and recovery time) correlate among them (Fig. S14).

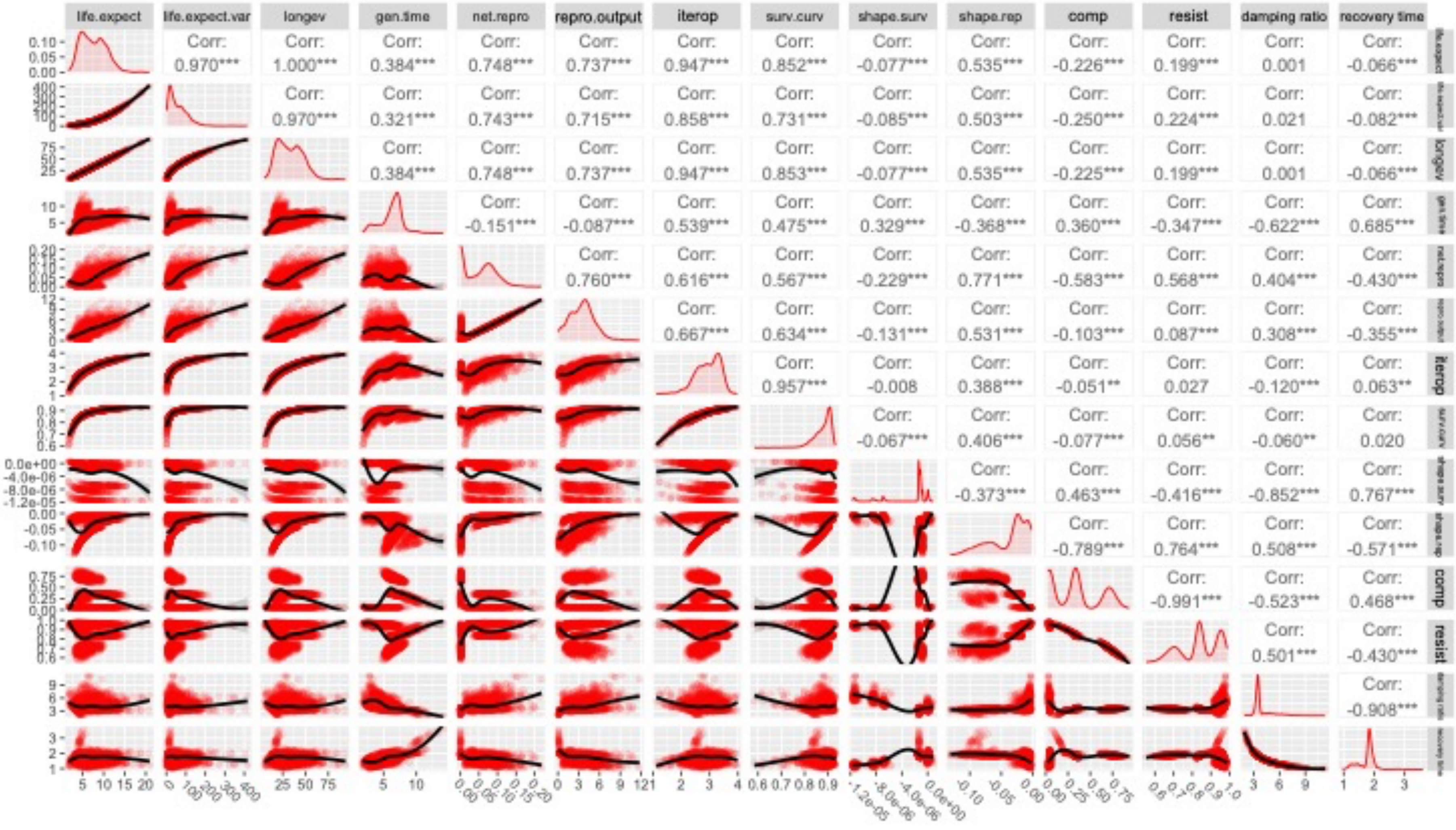


**Fig. S14.** Correlations and relationships among life history traits and demographic resilience components (resistance, compensation, damping ratio, and recovery time) estimated by stochastic Integral Projection Models of three species of lizards (*Copeoglossum nigropunctatum*, *Micrablepharus atticolus*, and *Tropidurus itambere*) from the Brazilian Cerrado savannas. *comp* = compensation; *gen.time* = generation time; *iterop* = degree of iteroparity; *life.expect* = life expectancy; *life.expect.var* = variance of life expectancy; *longev* = longevity; *net.repro* = net reproductive output expected lifetime production of offspring that start life by an individual also starting life; *repro.output* = net reproductive output calculated as the per-generation population growth rate; *resist* = resistance; shape*.rep* = 'shape' value of distribution of reproduction over size; *shape.surv* = 'shape' value of survival lifespan inequality; *surv.curv* = shape of the age-specific survivorship curve.

We also show how reproductive output and generation time vary among species and fire regimes of varying fire severity (Fig. S15). We can observe that the main differences among fire regimes derive from the reproductive output in *C. nigropunctatum*, generation time in *M. atticolus*, and both reproductive output and generation time in *T. itambere* (Fig. S15).

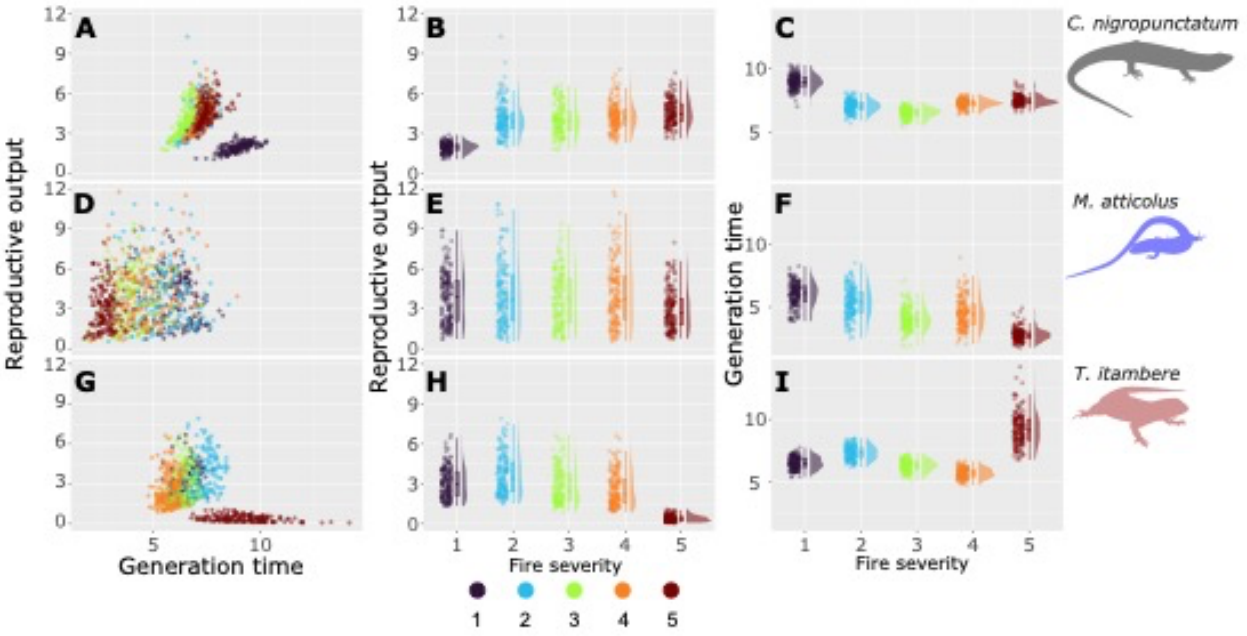


**Fig. S15.** Variation of reproductive output and generation time of populations of three lizard species (*Copeoglossum nigropunctatum*, *Micrablepharus atticolus*, and *Tropidurus itambere*) from the Brazilian Cerrado savannas in fire regimes of varying severity.
